# The Mitochondrial F-box protein 1 and DJ-1 homolog HSP31 support cellular proteostasis during mitochondrial protein import clogging

**DOI:** 10.1101/2025.10.13.682092

**Authors:** Gargi Mishra, Xiaowen Wang, Eamon Fitzpatrick, Auyon Ghosh, Xin Jie Chen

## Abstract

Mitochondrial biogenesis requires the import of ∼1,000-1,500 nuclear-encoded proteins across the Translocase of Outer Membrane (TOM) and the Translocase of Inner Membrane (TIM) 22 or 23 complexes. Protein import defects cannot only impair mitochondrial respiration but also cause mitochondrial Precursor Overaccumulation Stress (mPOS) in the cytosol. Recent studies showed that specific mutations in the nuclear-encoded Adenine Nucleotide Translocase 1 (ANT1) cause musculoskeletal and neurological diseases by clogging TOM and TIM22 and inducing mPOS. Here, we found that overexpression of *MFB1*, encoding the mitochondrial F-box protein 1, suppresses cell growth defect caused by a clogger allele of *AAC2*, the yeast homolog of Ant1. Disruption of *MFB1* synergizes with a clogger allele of *aac2* to inhibit cell growth. This is accompanied by increased retention of mitochondrial proteins in the cytosol, suggesting exacerbated defect in mitochondrial protein import. Proximity-dependent biotin identification (BioID) suggested that Mfb1 interacts with several mitochondrial surface proteins including Tom22, a component of the TOM complex. Loss of *MFB1* under clogging conditions activates genes encoding cytosolic chaperones including *HSP31*. Interestingly, disruption of *HSP31* creates a synthetic lethality with protein import clogging under respiring conditions. We propose that Mfb1 functions to maintain mitochondrial protein import competency under clogging conditions, whereas Hsp31 plays an important role in protecting the cytosol against mPOS. Mutations in DJ-1, the human homolog of Hsp31, and mitochondria-associated F-box proteins (eg., Fbxo7) are known to cause early-onset Parkinson’s disease. Our work may help to better understand how these mutations affect cellular proteostasis and cause neurodegeneration.

## INTRODUCTION

Mitochondria are essential organelles that perform a wide range of functions, including ATP production, metabolite synthesis, stress signaling, programmed cell death, and immune signaling. Mitochondria possess their own genome, called mitochondrial DNA (mtDNA). While the mitochondrial genome encodes a small number of proteins, the vast majority (approximately 1,000 to 1,500) are encoded by the nuclear genome. These proteins are translated in the cytosol and imported into mitochondria through specialized translocase complexes, with assistance from chaperones, membrane receptors, and motor proteins located on the outer and inner membranes or in the aqueous subcompartments of the mitochondrial matrix and the intermembrane space (Wiedemann and Pfanner 2017). Defects in protein import adversely impact mitochondrial biogenesis and function. Additionally, when import is defective, unimported mitochondrial precursors can accumulate in the cytosol, triggering a form of proteostatic stress known as mitochondrial Precursor Overaccumulation stress (mPOS) (Wang and Chen 2015; Coyne and Chen 2019). Various forms of mitochondrial damage can indirectly affect protein import efficiency, which in turn causes mPOS and decreases cell viability. Cells respond to mPOS by activating the proteasomal machinery to contain the unimported proteins from accumulating, a mechanism termed Unfolded Protein Response activated by Mistargeting of proteins (UPRam) (Wrobel *et al*. 2015). In addition to genetic mutations that directly affect components of the core protein import machinery, recent studies have shown that mutations in mitochondrial cargo proteins, such as Adenine Nucleotide Translocase 1 (ANT1), can get the protein stuck on import channels, leading to mitochondrial dysfunction, mPOS, and cell death (Coyne *et al*. 2023b). Notably, these “clogging” alleles of ANT1 have been implicated in human mitochondrial disorders such as autosomal dominant Progressive External Ophthalmoplegia (adPEO) (Kaukonen *et al*. 2000; Palmieri *et al*. 2005).

To counteract import clogging, cells have evolved quality control mechanisms that detect and resolve stalled import events on the mitochondrial surface. These include the mitochondrial Compromised Protein import Response (mitoCPR) and the mitochondrial protein translocation-associated degradation pathway (mitoTAD) (Weidberg and Amon 2018; Martensson *et al*. 2019). In these anti-clogging pathways, AAA-ATPases extract clogged and ubiquitinated precursors from the outer membrane, targeting them for proteasomal degradation in the cytosol. It was previously unclear as to which E3 ligase enzymes are responsible for initially detecting stalled mitochondrial proteins to ubiquitinate them for degradation by the proteasome. A recent study suggested that Rsp5 may be one such E3 ligase since it was found to ubiquitinate mitochondrial preproteins as they were being translated and imported at the outer mitochondrial membrane (Schulte *et al*. 2023). However, the full compendium of proteins that maintain protein import efficiency or assist in clearing stalled precursors remains to be defined.

In this study, we identified *MFB1* as a suppressor gene of mitochondrial protein import clogging in a genetic screen in *Saccharomyces cerevisiae*. *MFB1* encodes a mitochondrial F-box protein. Mfb1 has previously been shown to localize to the mitochondrial surface and play roles in maintaining mitochondrial morphology and the yeast replicative lifespan (Kondo-Okamoto *et al*. 2006; Pernice *et al*. 2016). It contains an F-box motif at its N-terminus that is believed to interact with members of a multi-subunit Skp1-Cullin-Fbox (SCF) E3 ligase complex. In a SCF complex, the F-box containing protein acts as a scaffold to which substrate proteins can bind, conferring specificity in ubiquitinating the proteins destined for degradation (Nguyen and Busino 2020). We found that Mfb1 is required for cell growth under mitochondrial protein import clogging conditions, likely via a SCF-independent mechanism. Strikingly, in the absence of Mfb1, cells activated expression of cytosolic stress-responsive genes, including the chaperone Hsp31. The Hsp31 family of proteins are homologous to human DJ-1, mutations of which are associated with juvenile Parkinsonism (Hague *et al*. 2003). Our data suggest that Mfb1 may maintain mitochondrial protein import competency, and that Hsp31/DJ-1 proteins provide a downstream defense line in the cytosol against mPOS.

## RESULTS

### MFB1 overexpression promotes cell growth under conditions of mitochondrial protein import clogging and defects in the protein import process

To investigate mechanisms that counteract mitochondrial import stress, we used a yeast model in which expression of a “pathogenic” *aac2* mutant inhibits cell growth. *AAC2* encodes the major isoform of adenine nucleotide translocase in aerobically grown yeast cells. The Aac2 protein is one of the most abundant mitochondrial proteins involved in ATP/ADP exchange across the inner membrane. Recent studies have shown that the A128P mutation of *AAC2*, equivalent to the pathogenic A114P allele in human ANT1 that causes autosomal dominant Progressive External Ophthalmoplegia (adPEO) (Kaukonen *et al*. 1999), increases the retention of the mutant protein at the TOM protein translocase complex on the outer membrane as well as at the TIM22 complex on the inner membrane (Coyne *et al*. 2023b). This causes a partial clogging of mitochondrial protein import. The growth of yeast cells expressing the *aac2^A128P^* allele from the *GAL10* promoter is strongly inhibited on galactose medium (Figure 1B). In a multicopy suppressor screen, we isolated two overlapping genomic clones that restored growth in cells expressing *GAL10-aac2^A128P^*. Sequence analysis revealed that *MFB1* is the only open reading frame present in the overlapping region between the two suppressor clones (Figure 1A). Further, we found that a subcloned version of full-length *MFB1* was sufficient to suppress the cell growth defect (Supplemental Figure S1). Multicopy expression of *MFB1* is therefore responsible for suppressing the growth defect of the cells with clogged mitochondrial protein import.

**Figure 1.**
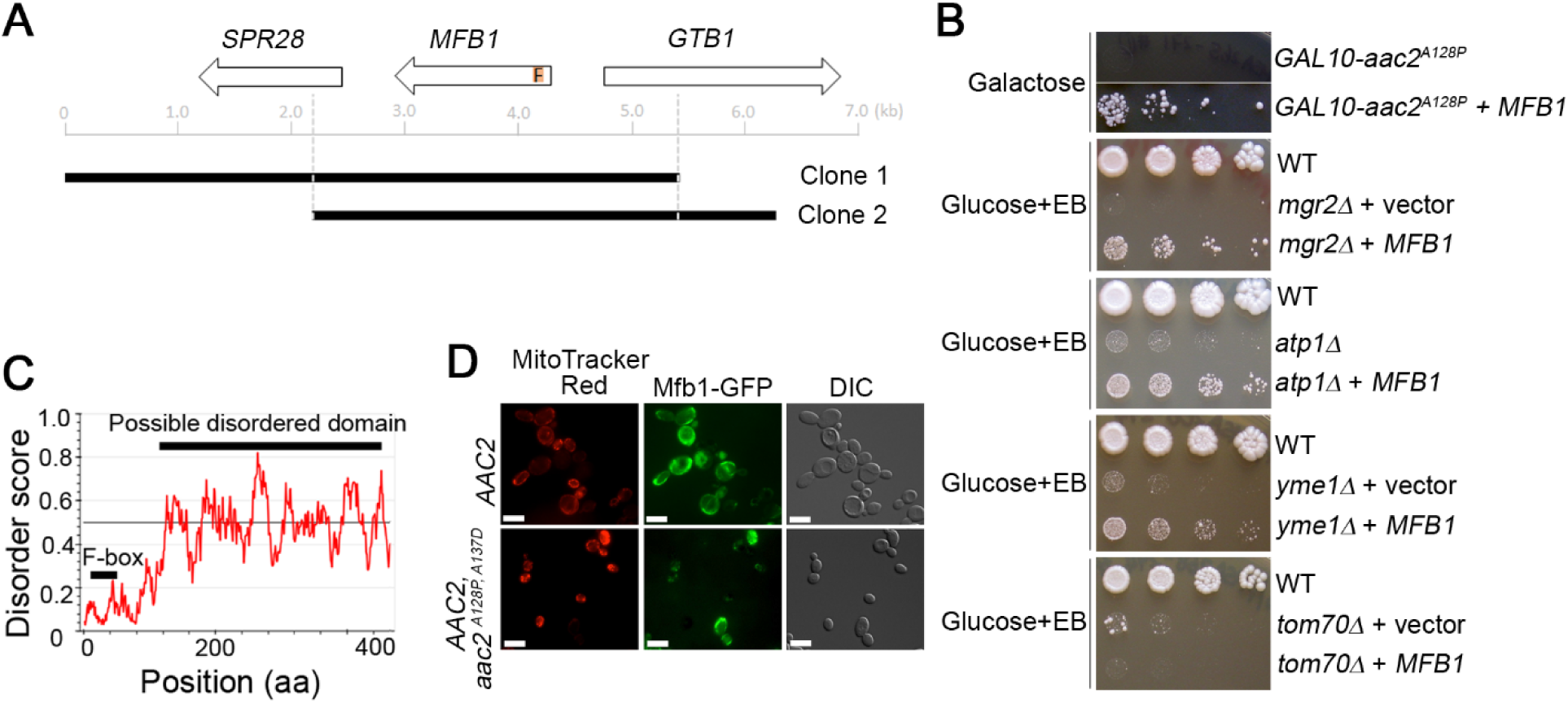
*MFB1* overexpression rescues cell growth under a variety of mitochondrial protein import defects. **(A)** *MFB1* on a multicopy plasmid rescues cell growth in *GAL10-aac2^A128P^* expressing cells, or in ethidium bromide (EB)-treated *mgr2Δ*, *atp1Δ, yme1Δ* but not *tom70Δ* cells. These mutant cells are ρ°-lethal due to low membrane potential and direct effect on protein import. EB eliminates mtDNA and further decreases membrane potential, leading to cell lethality. **(B)** Physical map of genomic clones on two multicopy plasmids having only *MFB1* as a shared gene that suppresses cell-lethality induced by *GAL10-aac2^A128P^*. **(C)** Mfb1 contains an F-box domain at the N-terminus and a disordered region at the C-terminus as predicted by IUPred2. **(D)** Fluorescence microscopy showing Mfb1-GFP localizing to mitochondria in both wild-type and the *aac2^A128P, A137D^* clogger strains. Mitochondria were stained by MitoTracker Red dye. Scale bar, 10 μm.

We subsequently found that *MFB1* overexpression can suppress various mutations that affect mitochondrial protein import. Mgr2 is a subunit of the TIM23 protein translocase (Matta *et al*. 2020) and *mgr2Δ* cells cannot tolerate the elimination of mtDNA that reduces membrane potential required for efficient protein import (Dunn *et al*. 2006), a phenotype known as ρ°-lethality. *ATP1* encodes the α-subunit of F1-ATPase. In addition to its role in ATP synthesis, F1-ATPase is also required for the maintenance of mitochondrial membrane potential (and therefore protein import) in ρ° cells (Chen and Clark-Walker 2000). As such, *atp1Δ* cells are ρ°-lethal (Chen and Clark-Walker 1999). Yme1 is a chaperone/protease on the mitochondrial inner membrane that is involved in protein quality control and membrane potential maintenance. Loss of Yme1 is also ρ°-lethal (Kominsky *et al*. 2002). We found that *MFB1* overexpression from a multicopy vector was able to suppress the ρ°-lethal phenotype associated with *mgr2Δ*, *atp1Δ*, and *yme1Δ* cells (Figure 1B). These findings suggest that *MFB1* may play a broader role in supporting mitochondrial protein import or in enhancing cell survival under protein import-compromised conditions. Disruption of *TOM70*, encoding a component of the TOM complex that tethers cytosolic chaperones to the outer mitochondrial membrane (Backes *et al*. 2021), is also known to cause ρ°-lethality (Dunn *et al*. 2006). Notably, *MFB1* overexpression failed to suppress the ρ⁰-lethal phenotype of *tom70Δ* cells, suggesting that Mfb1 function may depend on *TOM70* or a related docking site on the mitochondrial surface.

Mfb1 is composed of 465 amino acids. It is predicted to have an F-box domain on its N-terminus, followed by a largely disordered domain on its C-terminus (Figure 1C). Many F-box proteins are components of the SKP1–Cullin1–F-box protein (SCF) ubiquitin ligase complex, acting as substrate-recognition subunits that target specific proteins for degradation via the ubiquitin-proteasome pathway. However, some F-box proteins, including the mitochondria-associated Fbxo7 protein in humans are also proposed to be involved in various cellular functions without being part of the SCF complex (Nelson *et al*. 2013; Teixeira *et al*. 2016). Mfb1 has previously been shown to be involved in the maintenance of normal mitochondrial morphology (Durr *et al*. 2006; Schulte *et al*. 2023). It also plays a role in anchoring healthier-functioning mitochondria at the mother cell’s distal tip throughout the cell cycle and at the bud tip prior to cytokinesis (Kondo-Okamoto *et al*. 2006). Consistent with a previous report (Kondo-Okamoto *et al*. 2008), Mfb1 co-localizes with mitochondria in wild-type cells. Furthermore, Mfb1’s mitochondrial localization is not affected even in *aac2^A128P, A137D^*cells that display a severe clogging of protein import (Coyne *et al*. 2023b) (Figure 1D).

### Disruption of *MFB1* synergizes with mitochondrial protein import clogging to further inhibit cell growth and destabilize mtDNA

The suppression of the *aac2^A128P^* clogging allele and *mgr2Δ* by *MFB1* overexpression strongly supports the idea that *MFB1* promotes mitochondrial protein import competency. To further test this, we examined whether loss of *MFB1* synergizes with mitochondrial protein import stress to exacerbate defects in cell growth. As shown in Figure 2A, expression of *aac2^A128P^* causes a dominant cold-sensitive phenotype, especially at 15°C. The *mfb1Δ* mutant also exhibited mildly reduced growth at 15°C compared to 25°C and 30°C. Importantly, the *aac2^A128P^ mfb1Δ* double mutant showed markedly impaired growth at 15°C, beyond that seen in either single mutant. When single cells were spotted on YPD medium, 48.8% of *aac2^A128P^ mfb1Δ* cells either failed to form colonies or formed microcolonies that could not continue proliferating, compared to 14.4% and 2.2% for *aac2^A128P^* and *mfb1Δ* single mutants, respectively (Figure 2B and 2C). These data support a role for Mfb1 in promoting cell growth under conditions of mitochondrial protein import clogging. Since this growth defect was observed on glucose medium, where respiration is dispensable, the data suggest that Mfb1 enhances cell viability under import stress independent of mitochondrial respiration. We speculate that Mfb1 contributes to the maintenance of mitochondrial protein import competency during *aac2^A128P^*-induced clogging.

**Figure 2.**
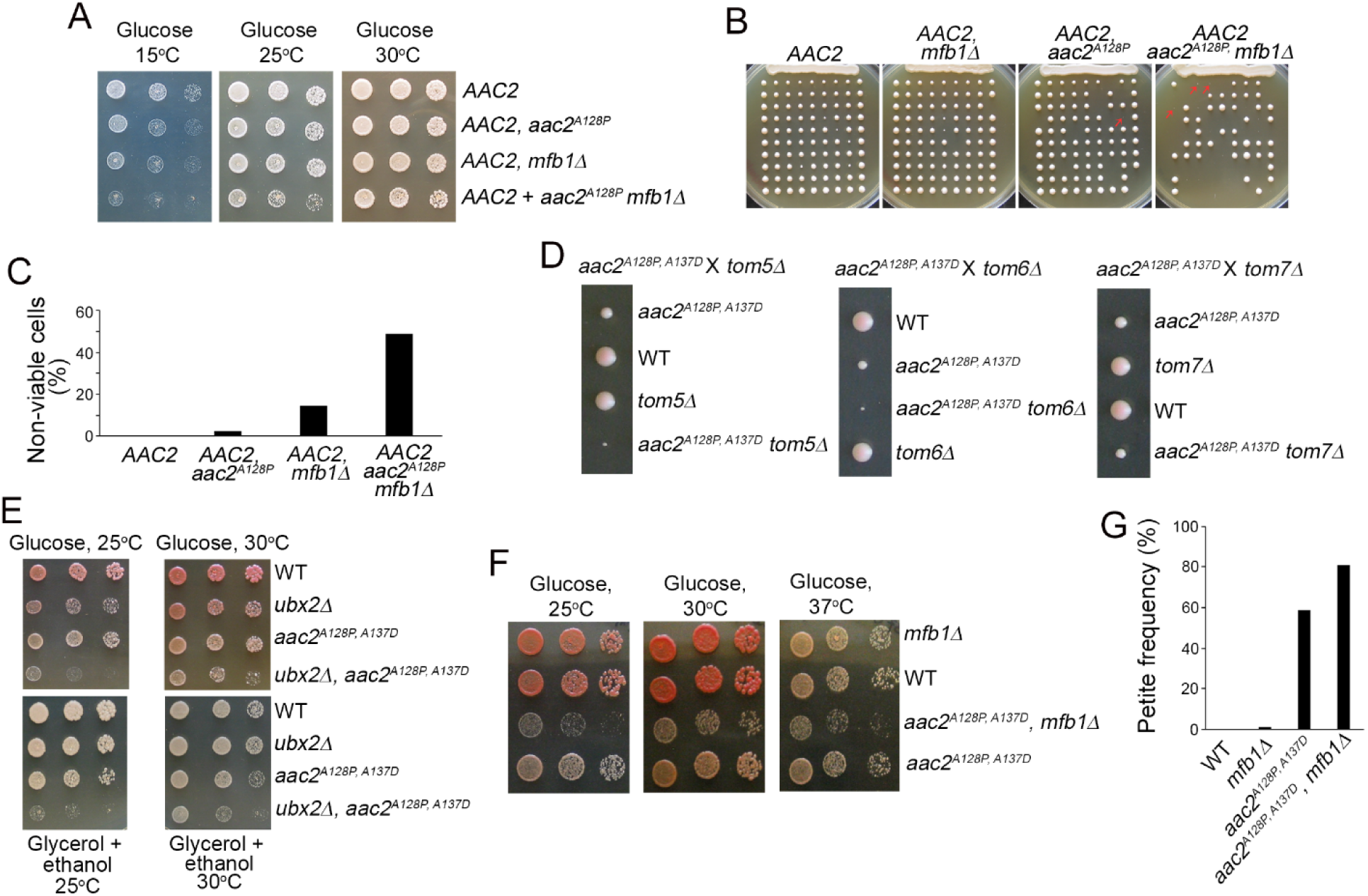
Disruption of *MFB1* under import clogging conditions worsens growth defects and decreases mtDNA stability. **(A)** Five-fold dilution series of yeast cells expressing the dominant single mutant *aac2^A128P^* clogger allele with or without concurrent *MFB1* deletion on a glucose medium. **(B)** Disruption of *MFB1* leads to either defective or non-viable microcolonies (red arrows) in *aac2^A128P^*background. Single cells from liquid YPD cultures at late-exponential phase were spotted onto a YPD agar plate by a Singer dissection microscope. **(C)** Quantification of non-viable cells/colonies shown in (b). **(D)** Disruption of *TOM5, 6* and *7* reduces the growth of clogger *aac2^A128P, A137D^* cells. A representative tetrad from each cross was dissected on synthetic complete glucose medium. **(E)** Disruption of *UBX2* reduces the growth of the clogger *aac2^A128P, A137D^*cells on fermentable (glucose) or non-fermentable (glycerol plus ethanol) media. **(F)** Disruption of *MFB1* reduces the growth of the clogger *aac2^A128P, A137D^* cells on synthetic complete glucose medium. (**G**) Disruption of *MFB1* increases the formation of petite colonies in the *aac2^A128P, A137D^*cells. The strains were grown overnight in complete glycerol plus ethanol medium, before being diluted in water and plated on YPD. All the strains are isogenic and harbor the *ade2* mutation which allows the development of red pigment if cells are respiratory competent. The petites were scored based on the development of white colonies due to mutations or loss of mtDNA. Approximately 300 colonies were counted for each strain.

The Aac2^A128P^ mutant protein impairs import along the full translocation pathway, from the outer to the inner mitochondrial membrane. In contrast, the double mutant Aac2^A128P, A137D^ preferentially blocks import at the Tom40 channel on the outer membrane (Coyne *et al*. 2023b). In tetrad dissections, the *aac2^A128P, A137D^* spores showed clearly reduced growth on glucose medium compared to wild-type cells. Supporting the outer membrane-specific import clogging model, *aac2^A128P, A137D^* growth was further impaired when non-essential components of the TOM40 complex Tom5 and Tom6, and to a lesser extent Tom7, were deleted (Figure 2D). Additionally, loss of *UBX2*, which helps cells tolerate outer membrane import clogging (Martensson *et al*. 2019), was synthetically lethal with *aac2^A128P, A137D^* at 25°C on both fermentable and non-fermentable carbon sources (Figure 2E).

Most notably, *MFB1* deletion also severely impaired the growth of *aac2^A128P, A137D^* cells on glucose medium, especially at 25°C (Figure 2F), reinforcing a role for Mfb1 in mitigating import clogging. Furthermore, *aac2^A128P, A137D^* cells frequently formed petite colonies, a hallmark of mtDNA instability. This petite frequency increased even further in *aac2^A128P, A137D^ mfb1Δ* cells (Figure 2G). These findings suggest that clogging caused by Aac2^A128P, A137D^ disrupts pathways required for mtDNA maintenance, either inside or outside the mitochondria. The additional loss of Mfb1 may intensify import stress, further destabilizing mtDNA and increasing petite formation. Altogether, our genetic data support a role for Mfb1 in preserving mitochondrial protein import and mtDNA stability under stress conditions.

### MiniTurboID studies revealed Mfb1’s proximity to the TOM complex and enrichment near inter-organelle contact sites

To better understand the functional relevance of Mfb1 at the mitochondrial surface, we used a proximity labeling approach to identify proteins in its immediate vicinity. We fused a miniTurbo biotin ligase (Branon *et al*. 2018) to the C-terminus of Mfb1 at the chromosomal locus in both *AAC2* wild-type and *aac2^A128P, A137D^* (“clogger”) strains. The Mfb1-miniTurbo fusion retained its normal mitochondrial localization (Supplemental Figure S2). Given the high-sensitivity nature of the assay, we performed three independent replicates in the wild-type and clogger (*aac2^A128P, A137D^*) cells expressing Mfb1-miniTurbo (WTmT and DMmT respectively). Consistent with its proximity to mitochondria, the assay revealed numerous mitochondrial proteins that may be proximal to Mfb1 (Figure 3A and 3B). The total number of Mfb1’s potential interactors appeared to be similar between the wild-type and *aac2^A128P, A137D^* cells. We found 15 and 14 potential interactors that appeared in all the three replicates of WTmT and DMmT cells (Supplemental Figure S3). Only four proteins, namely Tom22, Pet10, Ycp4, and Yet3, were consistently identified in all six replicate experiments (three WTmT and three DMmT), strongly suggesting that they reside in stable proximity to Mfb1 (Figure 3C).

**Figure 3.**
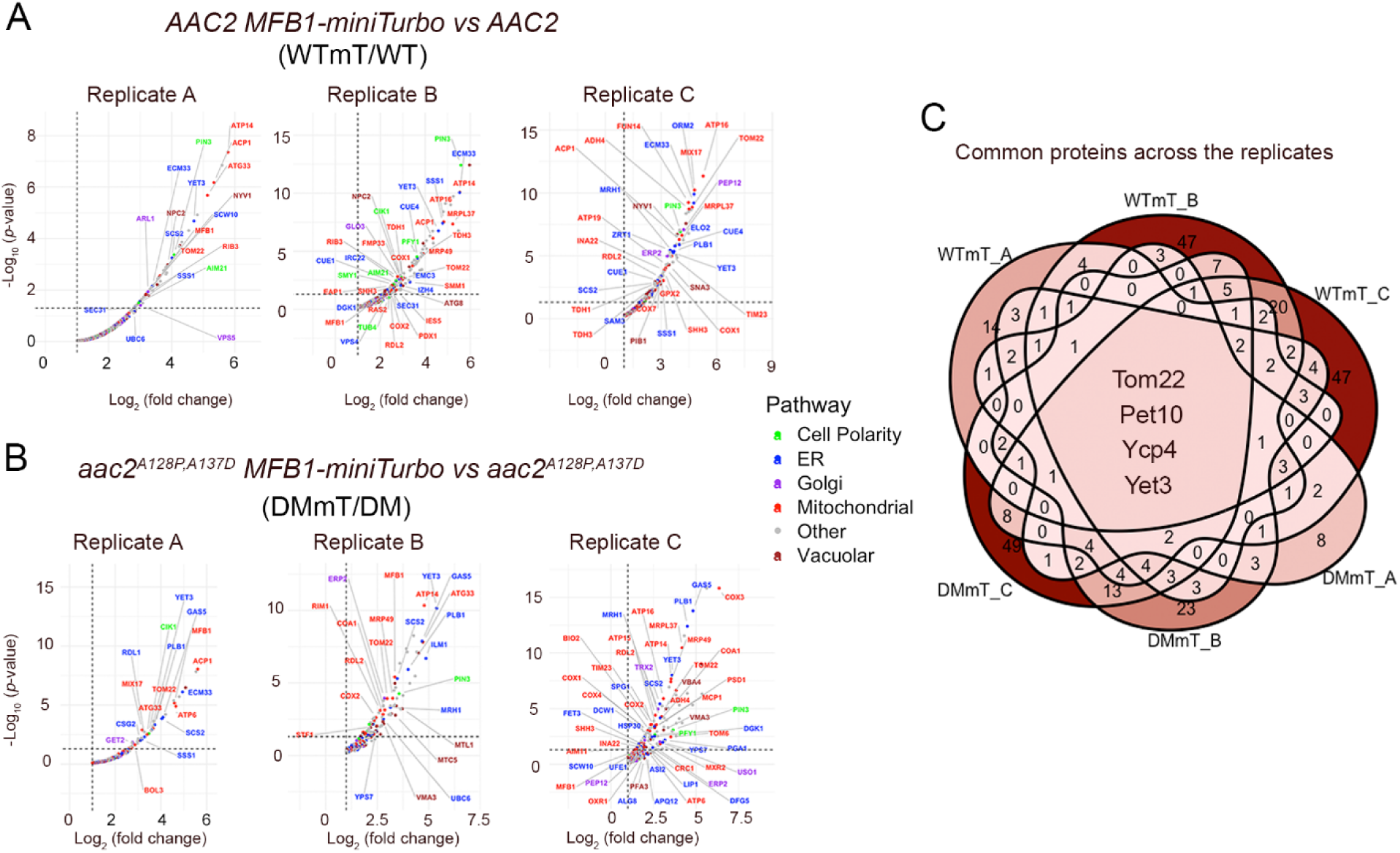
Mfb1 interactome analysis reveals its presence on the mitochondrial outer membrane in the proximity of interorganellar contact sites. **(A)** Enrichment of biotinylated proteins in the Mfb1-miniTurboID tagged cells in the background of wild type *AAC2*, normalized to untagged wild type. **(B)** Enrichment of biotinylated proteins in the Mfb1-miniTurboID tagged strains in the clogger cells (*aac2^A128P, A137D^*), normalized to untagged clogger. **(C)** Biotinylated proteins common to all replicates (wild type and clogger replicates combined).

Tom22 is a component of the TOM complex on the outer mitochondrial membrane, not only acting as a receptor for precursor proteins that have N-terminal mitochondrial targeting signals but also playing a crucial function in maintaining the integrity of the entire TOM complex (Dekker *et al*. 1998). Pet10 (or PLN1) is a lipid droplet protein mediating interaction with mitochondria through interactions with outer mitochondrial proteins including Tom22 (Pu *et al*. 2011; Gao *et al*. 2017). Yet3 is orthologous to the mammalian endoplasmic reticulum (ER) membrane protein BAP31 involved in ER-mitochondria interaction via Tom40 (Namba 2019). Ycp4 is a flavoprotein-like protein associated with mitochondria, but its physiological function is undefined (Reinders *et al*. 2006). Together with a previous study showing that Mfb1 co-immunoprecipitates with TOM71 (Kondo-Okamoto *et al*. 2008), our data further support the idea that Mfb1 may be peripherally associated with the outer mitochondrial membrane and in close proximity to the TOM complex. In support of this, Tom6, which is another TOM subunit, was also identified as a potential interactor in the *aac2^A128P, A137D^* background (Figure 3B). Mitochondrial matrix proteins were also detected, likely because they transiently pass near Mfb1 and the TOM complex during translocation across the outer membrane.

In addition to mitochondrial proteins, the proximity assay also identified numerous candidates from other organelles, including the ER (Cue1, Sec31, Ubc6), Golgi (Erp2, Trx2, Glo3), vacuole (Npc2, Nyv1, Vma3), and cytoskeleton (Pin3, Tub4, Smy1) (Figure 3, Supplemental Figure S3). Mfb1’s proximity to ER and Golgi proteins is again consistent with its presence on the outer mitochondrial surface and suggests that it may be enriched at inter-organelle contact sites. Finally, Mfb1’s proximity to cytoskeletal proteins supports its known role in mitochondrial anchorage during budding, a process important for maintaining the yeast replicative lifespan (Pernice *et al*. 2016).

### Mfb1 loss under clogging conditions leads to increased accumulation of mitochondrial proteins in the cytosol

Given Mfb1’s ability to suppress mPOS-mediated growth defects and its proximity to the import machinery, we hypothesized that Mfb1 promotes mitochondrial protein import competency. To test this, we cultured wild-type and *aac2^A128P, A137D^* (clogger) cells with or without *MFB1* in non-fermentable medium, where the demand for mitochondrial biogenesis is high. We then isolated post-mitochondrial cytosolic fractions and analyzed them using tandem mass-tagged (TMT) mass spectrometry to assess the accumulation of unimported mitochondrial proteins in the cytosol.

In *aac2^A128P, A137D^* cells, most mitochondrial proteins were reduced in the cytosol, with only a few showing increased levels (Figure 4A). This could indicate that the cytosolic proteostatic network can manage mPOS under respiratory conditions. Alternatively, reduced import efficiency and mtDNA instability may activate retrograde signaling pathways that repress mitochondrial biogenesis, thereby decreasing precursor protein synthesis. In contrast, the absence of Mfb1 in *aac2^A128P, A137D^* cells led to increased accumulation of many mitochondrial proteins in the cytosol (Figure 4B, 4C), consistent with the exacerbation of mPOS in the cytosol. Notably, Tom20, which is a core component of the import machinery, was among the enriched proteins. This may reflect impaired mitochondrial targeting of Tom20 itself, or a destabilization of the TOM complex upon loss of Mfb1 under clogging conditions. These findings support a model in which Mfb1 safeguards mitochondrial protein import by limiting precursor accumulation in the cytosol and/or maintaining TOM complex integrity.

**Figure 4.**
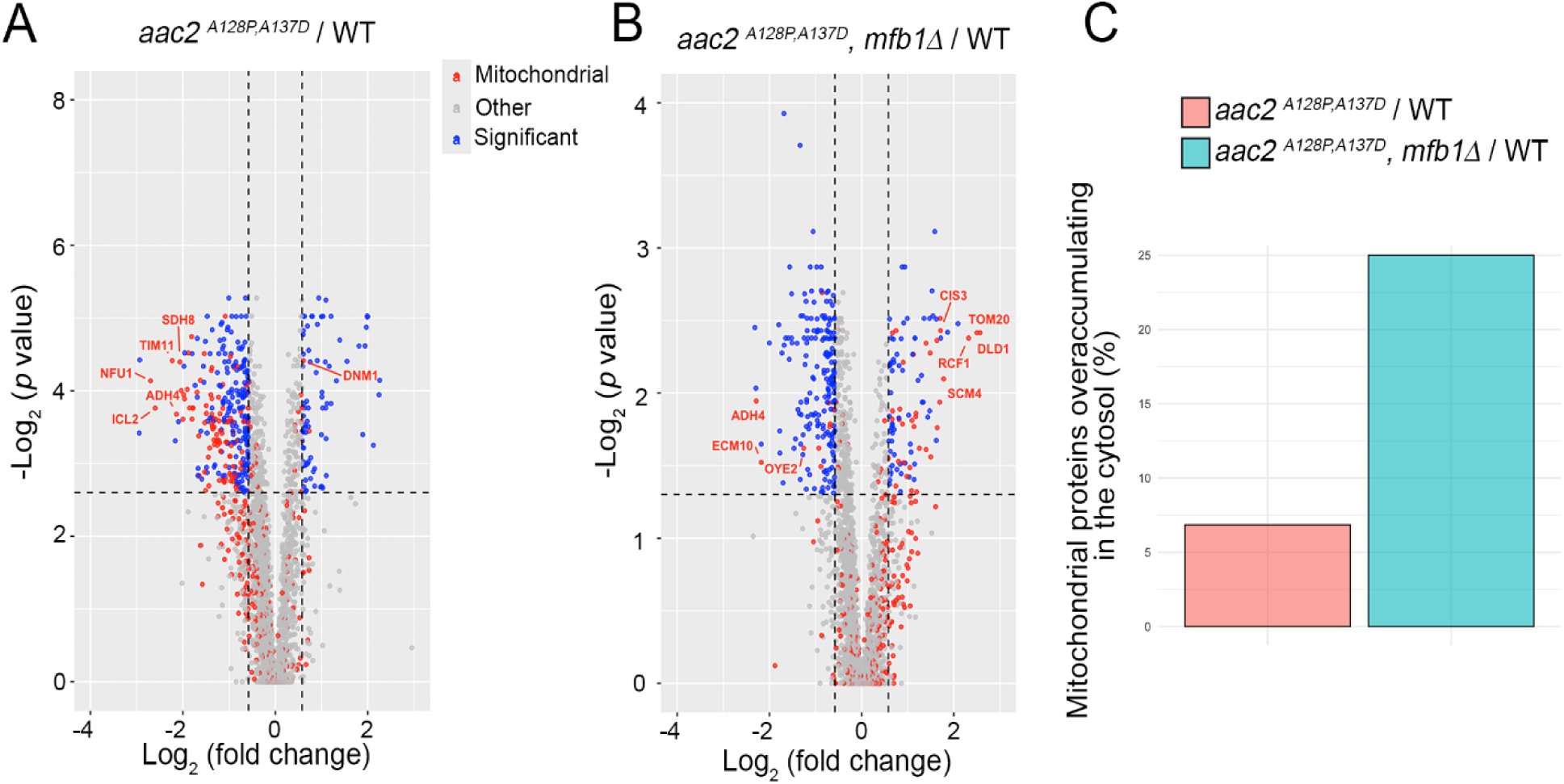
Disruption of *MFB1* increases the cytosolic retention of mitochondrial proteins under protein import clogging conditions as revealed by TMT-mass spectrometry analysis of cytosolic fractions. **(A)** Mitochondrial proteins enriched or depleted in clogger cells (*aac2^A128P, A137D^*) relative to wild type when grown under non-fermentable conditions. **(B)** Mitochondrial proteins enriched or depleted in clogger cells lacking *MFB1* (*aac2^A128P, A137D^ mfb1Δ*) relative to wild type. **(C)** Percentage of mitochondrial proteins enriched relative to all significantly enriched proteins in *aac2^A128P, A137D^* (pink) or *aac2^A128P, A137D^ mfb1Δ* (blue) cells.

### Disruption of *MFB1* activates proteostatic stress responses under mitochondrial protein import clogging conditions

To examine whether Mfb1 influences transcriptional adaptation in the setting of mitochondrial protein import clogging, we performed RNA-seq on wild-type, *mfb1Δ*, *aac2^A128P, A137D^* (DM), and *mfb1Δ aac2^A128, A137D^* (DM *mfb1Δ*) strains grown under respiratory conditions. Pairwise comparisons revealed that the clogger (DM) allele by itself strongly induced iron homeostasis genes (e.g., *FET3, FRE1, FRE6, FIT3*) (Figure 5A, Supplemental Table 1), which is a hallmark of mutants with severe mitochondrial damage (Veatch *et al*. 2009). Mitochondrial protein import clogging broadly repressed genes encoding ribosomal (e.g., *RPL34A, RPL18A, RPS16B, RPS13*) components, suggesting reduction of protein synthesis as an adaptation to heightened proteostatic stress (Supplemental Figure S4A). Several proteasomal components (*PRE6-9*) were also repressed, suggesting remodeling of proteosomal function. Loss of Mfb1 further amplified this transcriptional remodeling. For example, in the DM *mfb1Δ* relative to *mfb1Δ* comparison, stress response programs were expanded to include known mitochondrial quality control factors (*YME1, YTA12, MSP1, and RSP5*), autophagic and mitophagic factors (*ATG5, ATG11, ATG32*) (Figure 5B), while genes involved in protein synthesis were repressed (Supplemental Figure S4B).

**Figure 5.**
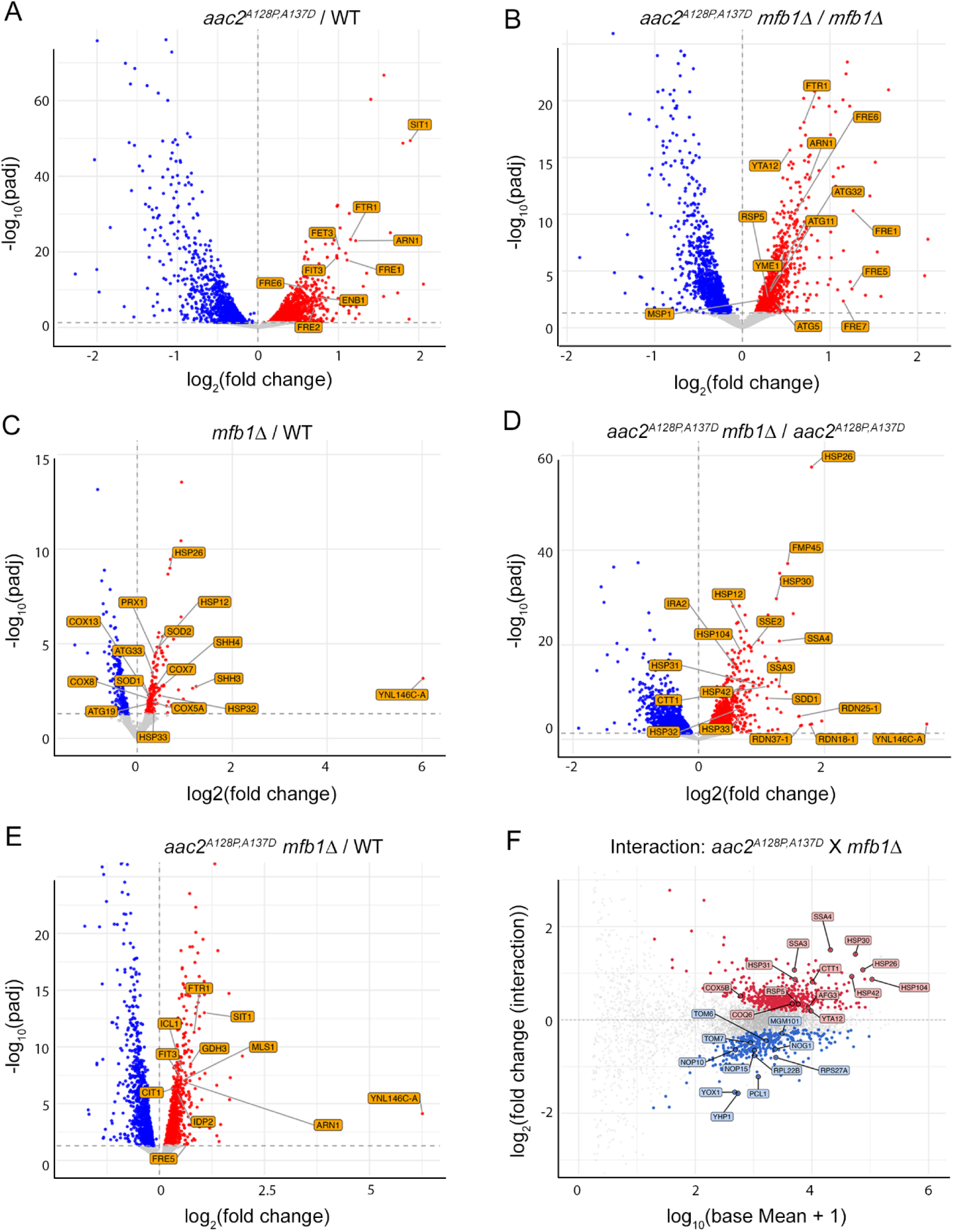
Transcriptomic analysis reveals the activation of unique cellular pathways under conditions of protein import clogging and *MFB1* loss. **(A)** Volcano plot labeling a subset of genes involved in iron homeostasis that are activated under clogging (*aac2^A128P, A137D^*) conditions relative to wild type when *MFB1* is present. (**B**) In addition to iron homeostasis, a subset of mitochondrial quality control genes become activated during clogging conditions relative to wild type when MFB1 is absent. (**C**) *MFB1* loss relative wild type leads to upregulation of small chaperone proteins in the cytosol. (**D**) *MFB1* loss relative to wild type leads to a drastic upregulation of many different chaperone protein families during protein import clogging conditions. (**E**) Clogging with concurrent *MFB1* loss compared to wild type reveals the upregulation of genes involved in TCA cycle and peroxisomal function, both of which are both hallmarks of an activated mitochondrial retrograde signaling response. (**F**) An MA (Minus Average)-plot showing the genes with the strongest interaction across genotypes in the multifactorial analysis. The y-axis shows the difference, i.e. log normalized fold change resulting from the interaction between *mfb1Δ* and *aac2^A128P, A137D^*. The x-axis indicates the log-normalized average expression level across all samples for every given gene. Genes labeled in red are more upregulated when comparing in *mfb1Δ* vs *MFB1* wild type under import clogging conditions relative to wild type. Similarly, genes labeled in blue are more downregulated when comparing *mfb1Δ* vs *MFB1* wild type under import clogging conditions relative to wild type.

*MFB1* deletion alone was sufficient to upregulate small heat shock protein genes (*HSP12, HSP26* as well as *HSP32* and *HSP33*), redox regulators (*PRX1, SOD1, SOD2*), and oxidative phosphorylation subunits (e.g., *COX5A, COX8, COX13*) (Figure 5C), while repressing members of the telomere maintenance gene family *YRF1* (Supplemental Figure S4C), indicating a basal role for Mfb1 in maintaining cellular homeostasis. When clogging was introduced into this background allowing us to compare the DM *mfb1Δ* versus DM conditions, we saw a further increase in compensatory proteostatic pathways (Figure 5D, Supplemental Table 2) including the *HSP70* family members (*SSE2, SSA3 and SSA4*), cytosolic deaggregase (*HSP104*), as well as the upregulation of the multifaceted chaperone-coding gene *HSP31* which had not been upregulated in the *mfb1Δ* versus WT comparison. *HSP31* belongs to the same family as *HSP32* and *HSP33* (Miller-Fleming et al. 2014). However, it appears to have non-redundant roles compared to *HSP32* and *HSP33* (Bankapalli et al. 2015). Importantly, *HSP31* is the yeast ortholog for human DJ-1, mutations in which are linked to early-onset autosomal recessive forms of Parkinson’s disease (Abou-Sleiman *et al*. 2003; Bonifati *et al*. 2003; Hague *et al*. 2003). While Hsp31 has been shown to act as a responder to oxidative stress and for its glyoxylase activity to reduce mitochondrial damage (Bankapalli *et al*. 2015; Alshammari 2025), its physiological role as a molecular chaperone is still unclear.

Next, when comparing the most growth-defective *aac2^A128P, A137D^ mfb1Δ* to WT, we noticed an additive effect (Figure 5E, Supplemental Table 3). In addition to seeing the upregulation of the iron homeostasis genes that we had seen in clogging alone (Figure 5A), we also observed the upregulation of genes belonging to various metabolic pathways such as the citric acid cycle and the glutamate synthesis (eg., *CIT1, CIT2, CIT3, ICL1, IDP2 MLS, GDH3, PUT1 and IDP2*) as well as that of glucose transporters (eg., *HXT1, HXT5, HXT16*, and *HXT2*). Activation of the glyoxylate cycle in the peroxisome promotes anaplerotic reactions to supply acetyl-CoA and citrate to mitochondria (Kunze *et al*. 2006). Increased glutamate biosynthesis provides all the nitrogen used in various biosynthetic reactions (Walker and van der Donk 2016). Both these pathways are direct targets of a mitochondria-to-nucleus retrograde signaling response which is activated under conditions of severe mitochondrial damage (Liu and Butow 2006). Thus, loss of *MFB1* under clogging conditions further heightens cellular stress, which reconfigures anaplerotic reactions and increases carbohydrate scavenging.

Notably, when we compared *mfb1Δ* to wild type *MFB1* with or without the presence of clogging, we consistently observed the upregulation of an uncharacterized open reading frame *YNL146C-A* (Figure 5C, 5D, and 5E). We designated this gene as *AMK1* (*A*ctivated by *M*FB1 *K*nockout). Although *AMK1* deletion did not impair growth or genetically interact with *mfb1Δ*, *amk1Δ* cells modestly increased petite frequency in the clogger (DM) background (Supplemental Figure S5).

Finally, to further validate the genetic interactions, we applied a multifactorial model contrasting the effect of *MFB1* loss in WT versus in the clogger (DM) background. This identified ∼825 genes with significant interaction effects, many involved in stress response, protein quality control, and metabolic adaptation (Figure 5F, Supplemental Table 4). Together, these results indicate that the clogger mutant triggers an iron starvation response, while concomitant loss of Mfb1 markedly exacerbates cytosolic proteostatic stress and reconfigures cellular transcriptional programs to favor stress tolerance at the expense of growth.

### Disruption of *HSP31* is synthetically lethal with *aac2^A128P, A137D^* under respiring conditions

RNA-sequencing revealed that several cytosolic chaperones were upregulated in *mfb1Δ* cells under clogging conditions, suggesting that they contribute to cell survival during severe mitochondrial protein import stress. We focused on Hsp31, the yeast homolog of human DJ-1 due to its direct clinical relevance in contributing to Parkinson’s disease (Abou-Sleiman *et al*. 2003; Hague *et al*. 2003; Alshammari 2025). We found that *hsp31Δ*, like *mfb1Δ*, exacerbated the growth defect of *aac2^A128P, A137D^* cells at 25°C or 37°C, especially on non-fermentable carbon sources (Figure 6A). This effect was less pronounced at 30°C. Strikingly, *mfb1Δ hsp31Δ aac2^A128P, A137D^* cells failed to grow at 37°C. This supports the idea that *HSP31* upregulation in *mfb1Δ aac2^A128P, A137D^* cells (Figure 5D) is a specific protective response under severe clogging stress. Although *hsp31Δ* alone does not significantly increase mtDNA instability in the *aac2^A128P, A137D^* background, it markedly enhances petite colony formation when combined with *mfb1Δ* (Figure 6B).

**Figure 6.**
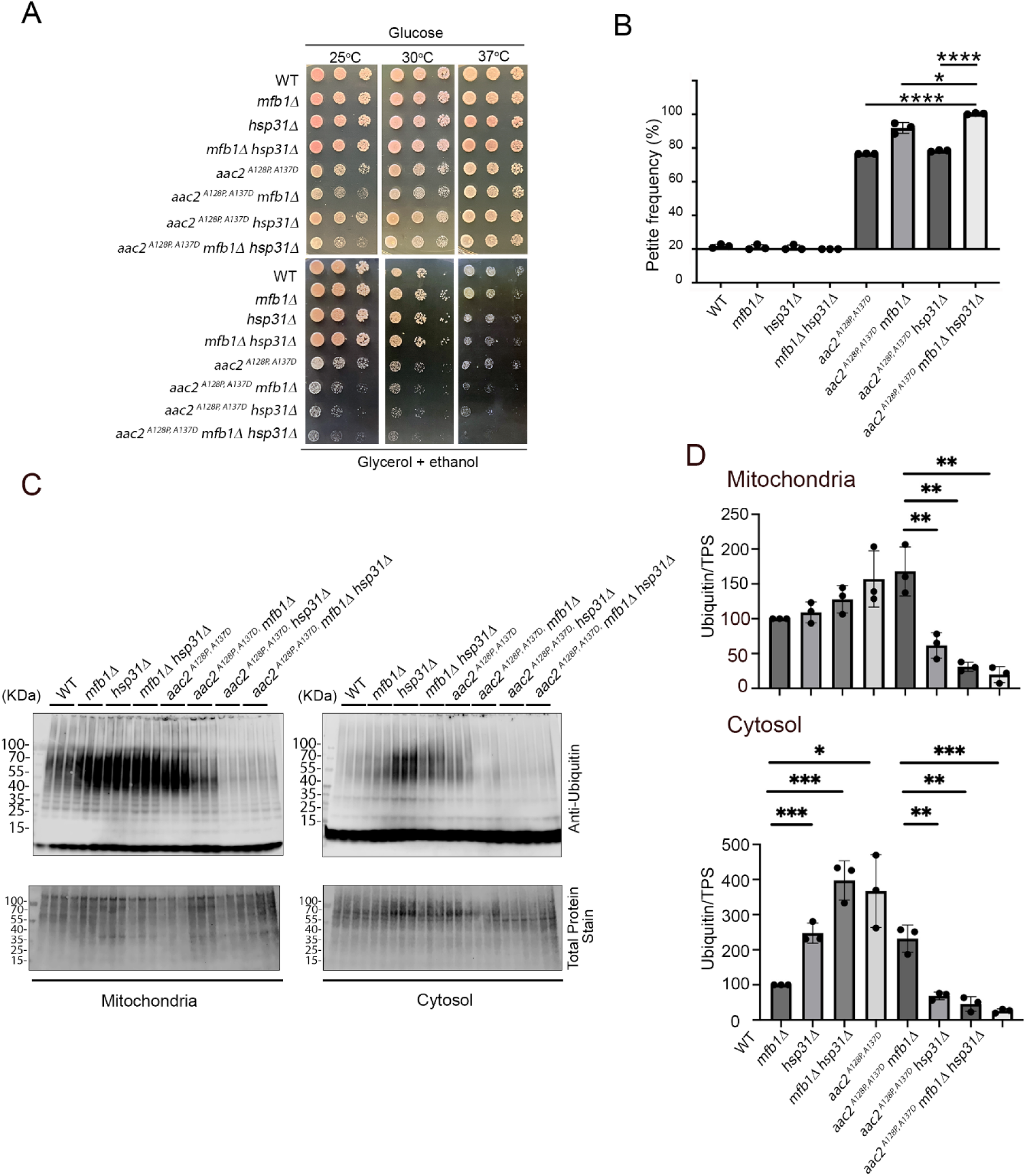
Disruption of *HSP31* creates a synthetic growth defect with *aac2^A128P, A137D^* and *aac2^A128P,A137D^ mfb1Δ* and increases the levels of ubiquitinated proteins in the cytosol under respiring conditions. **(A)** Five-fold dilution series for cells grown on fermentable YPD (glucose) after two days; or non-fermentable YPGE (glycerol and ethanol) medium after three days (25°C), or two days (30° and 37°C). **(B)** Petite frequency of yeast strains shown in (A). **(C)** SDS-PAGE showing protein ubiquitination detected in the mitochondrial and cytosolic fractions from strains as indicated. **(D)** Quantification of western blots shown in (C). Error bars represent the standard deviation of means with three independent biological replicates. *P* values were calculated using Student’s t test. *, p < 0.05; **, p < 0.01; ***, p < 0.001; ****, p < 0.0001.

Consistent with mitochondrial protein import clogging and the resulting mPOS in the cytosol, western blot analysis showed that both *hsp31Δ* and *mfb1Δ* increased protein ubiquitination in cytosolic but not mitochondrial fractions when compared with wild-type cells (Figure 6C and 6D). In contrast, cytosolic ubiquitination was reduced in *aac2^A128P, A137D^* mutants lacking *MFB1*, *HSP31*, or both, suggesting that the ubiquitination machinery may be impaired in these severely compromised cells. Importantly, protein ubiquitination was elevated in the Triton X-insoluble fraction of *aac2^A128P, A137D^* and *hsp31Δ* cells (Supplemental Figure S6), consistent with increased formation of insoluble protein aggregates. Under the various protein import stress conditions, Hsp31 remained diffusely localized throughout the cytosol (Supplemental Figure S7). Together, these findings support an intrinsic role for Hsp31 in maintaining cytosolic proteostasis during mitochondrial import stress, likely by stabilizing or facilitating degradation of unimported mitochondrial preproteins.

## DISCUSSION

In response to oxygen availability, cells upregulate mitochondrial biogenesis and increase the rate of mitochondrial protein import. Defective protein import can be detrimental. In addition to the potential impact on oxidative metabolism, it can also lead to the toxic accumulation of unimported mitochondrial proteins in the cytosol (Coyne and Chen 2018). Human adenine nucleotide translocase isoform 1 (ANT1) is primarily involved in ADP/ATP exchange across the inner mitochondrial membrane (Henderson and Lardy 1970). It belongs to the mitochondrial carrier protein family (Kunji *et al*. 2020), expressed mainly in the highly oxidative tissues including heart, skeletal muscle, and neurons. Missense mutations in *ANT1* cause dominant pathologies (Kaukonen *et al*. 1999; Kaukonen *et al*. 2000; Napoli *et al*. 2001; Komaki *et al*. 2002; Siciliano *et al*. 2003; Deschauer *et al*. 2005; Wang *et al*. 2008; Liu and Chen 2013; Thompson *et al*. 2016; Simoncini *et al*. 2017). It is well conserved across many species, including in yeast where its ortholog is called the ADP/ATP carrier protein 2 or *AAC2*. Genetic manipulation of the yeast *AAC2* has allowed for a meticulous dissection of the mechanisms that underlie human ANT1 pathologies (Mishra *et al*. 2023). Mutations in *AAC2* that are equivalent to the pathological alleles of *ANT1* not only affect nucleotide transport kinetics (Fontanesi *et al*. 2004), but also cause severe clogging of the mitochondrial protein import pathway, likely through aberrant interactions of the mutant protein with the protein import machinery (Coyne *et al*. 2023b). Import clogging subsequently leads to proteostatic stress in the cytosol (mPOS) that eventually overwhelms the cell’s capacity to maintain homeostasis (Wang and Chen 2015). In the current study, we used a yeast model to identify cellular pathways that promote cell survival under import clogging conditions. We found that the mitochondrial F-box protein Mfb1 and the cytosolic heat shock protein Hsp31 play an important role in the survival of respiring cells expressing clinically relevant alleles of *aac2* that clog mitochondrial protein import.

We identified *MFB1* from a genetic screen as a suppressor of *aac2^A128P^*, equivalent to the pathogenic *ant1*^A114P^ allele in humans. Most of the genetic suppressors identified in this screen were involved in maintaining cytosolic proteostasis (Wang and Chen 2015). *MFB1*, however, was unique in that it localizes to the mitochondrial surface, suggesting a more focused role in supporting the growth of cells clogged for mitochondrial protein import. Previously, Mfb1 has been characterized for its role in mitochondrial dynamics and the inheritance of mitochondria during cell division. For instance, disruption of *MFB1* has been shown to affect mitochondrial organization and connectivity (Kondo-Okamoto *et al*. 2006), the asymmetric inheritance of mitochondria in dividing cells (Durr *et al*. 2006; Pernice *et al*. 2016), and the replicative lifespan of yeast cells (Yang *et al*. 2022). Our findings add another pertinent layer to Mfb1’s function by showing that it also contributes to the maintenance of mitochondrial protein import competency. Specifically, we show that *MFB1* overexpression rescues cell growth under various mPOS-inducing conditions, while its deletion exacerbates precursor accumulation, impairs growth, and contributes to mtDNA instability. Since *MFB1* overexpression suppresses not only *aac2*-induced protein import clogging but also other types of stressors that affect protein import (e.g., *mgr2Δ*, and the ρ°-lethality phenotype of *atp1Δ* and *yme1Δ* cells), it is likely that Mfb1 plays a more general role in maintaining mitochondrial protein import competency. Since prior studies have established that Mfb1 tethers mitochondria at the mother cell tip throughout the cell cycle and at the bud tip of daughter cells during budding (Pernice *et al*. 2016; Yang *et al*. 2022), it is thus possible that Mfb1 tethers import-competent mitochondria to cytoskeletal structures, facilitating the transmission of healthy organelles to the daughter cells during aging. We propose that Mfb1 plays a role in the maintenance of mitochondrial protein import competency, which could have downstream effects on cellular processes such as mitochondrial dynamics. This is reminiscent of a recent study showing that loss of the mitochondrial morphodynamic protein Mitofusin-2 affects the integrity of the mitochondrial protein import machinery (Joaquim *et al*. 2025).

So how does Mfb1 contribute to the maintenance of mitochondrial protein import efficiency? Mfb1 is a yeast F-box protein (Durr *et al*. 2006; Kondo-Okamoto *et al*. 2006). Classically, F-box proteins act as part of a modular E3 ligase complex called Skp1-Cullin-F box or SCF complex. However, F-box proteins can have non-SCF functions as well (Nelson *et al*. 2013). Mfb1 seems to interact with Skp1, but not Cdc53, the yeast homolog for Cullin (Kondo-Okamoto *et al*. 2006). Skp1 is supposed to interact with F-box proteins through the F-box motif. Interestingly, the F-box motif of Mfb1 has been shown to be dispensable for the maintenance of mitochondrial morphology (Kondo-Okamoto *et al*. 2006). This implies that the mitochondrial morphodynamic function of Mfb1 may be independent of SCF complex interaction and protein ubiquitination. This is consistent with our finding that proximity-based biotin ligation followed by mass spectrometry analysis of isolated mitochondria failed to identify Skp1 and Cdc53 as Mfb1’s interactors. Notably, the Mfb1-Skp1 interaction was previously observed in a study where authors used whole cell lysates for immunoprecipitation (Kondo-Okamoto *et al*. 2006). It therefore remains possible that a smaller cytosolic fraction of Mfb1 may interact with Skp1, while the Mfb1 located on the mitochondrial surface does not.

Mfb1 has been shown to localize to the mitochondrial surface in a Tom70-dependent manner (Kondo-Okamoto *et al*. 2008). This suggests that Mfb1 is recruited to mitochondria either via direct interaction with Tom70, or via interaction with another TOM component with which the interaction is disrupted in the absence of Tom70. Consistent with this, we show that *MFB1* overexpression does not improve the growth of ρ° cells in the absence of Tom70, possibly due to a failure to localize to the mitochondrial surface.

Interestingly, our miniTurboID-MS experiment does not suggest a strong interaction between Mfb1 and Tom70. Instead, we identify several other mitochondrial outer membrane proteins in the vicinity of Mfb1. Among these proteins is Tom22, a precursor receptor on the TOM complex. These data provide further support for the idea that Mfb1 is localized in the vicinity of the TOM complex. Mfb1 may directly stabilize the TOM complex under clogging conditions, help unclog the protein translocation channel, and/or improve the microenvironment on the mitochondrial surface to facilitate preprotein targeting. Based on our TMT-MS study, the increased cytosolic accumulation of Tom20 in *aac2^A128P, A137D^ mfb1Δ* cells supports the idea that Mfb1 may help maintain TOM complex integrity during import stress. Altogether, these functions may help the maintenance of mitochondrial protein import efficiency and cell viability.

It is noteworthy that, in addition to Tom22, Pet10 and Yet3 were also identified as potential interactors of Mfb1. Pet10 is a lipid droplet protein mediating interaction with mitochondria through outer mitochondrial proteins including Tom22 (Pu *et al*. 2011; Gao *et al*. 2017). Yet3 is involved in ER-mitochondria interaction via Tom40 (Namba 2019). TOM components have been shown to serve as ER-mitochondria tethers (Ellenrieder *et al*. 2017), which may facilitate the import of precursor proteins that have been temporarily docked on the ER surface such as in the ER-SURF pathway (Koch *et al*. 2024). It is possible that Mfb1 localizes to regions engaged in active interactions with other organelles to facilitate inter-organellar communication including protein trafficking, or to sites of greater protein import on the OMM. The latter could be particularly relevant considering a recent study that describes how the mitochondrial membranes remodel to facilitate the co-translational import of mitochondrial proteins (Chang *et al*. 2025). The reduced distance observed between OMM and IMM around regions of co-translational protein import could contribute to changes in mitochondrial morphology. If Mfb1 maintains import competency by interacting with the TOM complex, its loss could destabilize regions with higher import frequency, subsequently leading to morphological changes.

Transcriptomic analysis provided further support for a role of Mfb1 in alleviating cellular stress under import clogging conditions. First, we employed a stratified analysis approach comparing WT, *mfb1Δ*, aac2^A128P,A137D^ (DM), and DM mfb1Δ strains grown under respiratory conditions. This strategy allowed us to identify differences between two genotypes at a time and to determine the effects of *mfb1Δ* and *aac2^A128P, A137D^* as genetic modifiers. It was interesting that in the *mfb1Δ* background, the additional introduction of import clogging led to an increase in genes that encode proteins which function on or near the mitochondrial surface (*MSP1, RSP5*) or within mitochondria (*YME1, YTA12*) (Figure 5B). However, in the *aac2^A128P, A137D^* (DM) background, the additional deletion of *MFB1* led to an increase in genes that encode a wide variety of cytosolic chaperones (Figure 5D). This suggests that under Mfb1-deficient conditions with newly induced clogging, the cell acts to rectify import stress by directly trying to triage/repair damaged mitochondria; while under clogging conditions, Mfb1 loss triggers a “proteostatic crisis” which requires the activation of cytosolic chaperones to fix. Finally, the stress of import clogging alone or concurrently with *MFB1* disruption, led to a downregulation of genes involved in ribosomal biogenesis, nucleolar function, and non-coding RNA processing (Supplemental Figures S4A, S4B, S4D, and S4E) which is consistent with reduced cytosolic translation and ribosomal function to alleviate proteostatic stress. Interestingly, Mfb1 loss alone relative to wild type led to a downregulation of the *YRF1* gene family members (Supplemental Figure S4C) which are known to be involved in telomere maintenance (Yamada *et al*. 1998). The physiological relevance of this transcriptomic finding is currently unclear.

Stratified analysis also allowed us to capture an additive interaction effect in our transcriptomic dataset. We found that Aac2^A128P, A137D^-induced clogging activates an iron starvation response. A similar response has been previously observed in ρ° cells that have reduced inner membrane potential and defective protein import (Veatch *et al*. 2009). Interestingly, clogging with concurrent *MFB1* loss leads to the activation of retrograde (RTG) signaling and increased glucose transport, in addition to the iron starvation response. RTG activation is suggestive of severe mitochondrial damage and drastic metabolic reprogramming, allowing cells to undergo anaplerosis and replenish glutamate stores (Liu and Butow 1999). These data suggest that as the severity of mitochondrial protein import stress increases, there is a corresponding “step-wise” increase in the types of cellular metabolic pathways that get dysregulated.

To be comprehensive in our understanding of the yeast cell’s transcriptional responses to import clogging and *MFB1* loss, we also employed a multifactorial approach to further validate the genetic interactions within all four yeast strains included in our study. This approach allowed us to define how *MFB1* loss affects gene expression differently based on whether the cell is undergoing protein import clogging or is in protein import homeostasis. The multifactorial design was able to improve our confidence in the results we obtained from the stratified analysis and help determine which genes are differentially expressed the most in context of the interaction between *mfb1Δ* and *aac2^A128P, A137D^*. For instance, *SSA4* is drastically upregulated when *MFB1* is deleted under clogging (*aac2^A128P, A137D^*) conditions but barely changed in expression when *MFB1* is deleted under wild type *AAC2* conditions. Other notable genes that experience the strongest differences in expression include *HSP30*, *HXT15*, *HSP31*, and *HSP26* among others (Supplemental Table 4).

While *HSP31* wasn’t the most upregulated gene in our dataset, we decided to focus our downstream validations on it for two primary reasons. The first reason was that Hsp31 is known for its role in oxidative stress defense, protein glyoxylation (Bankapalli *et al*. 2015), post-diauxic shift growth (Miller-Fleming *et al*. 2014), α-synuclein handling (Tsai *et al*. 2015; Aslam and Hazbun 2016), and more recently in redox-dependent mitochondrial dynamics (Biswas and D’Silva 2025), but its role as a molecular chaperone has not been well established. In our dataset, the entire Hsp31 gene family (*HSP31, HSP32, HSP33*) was upregulated. The proteins encoded by these genes are localized to the cytosol. However, during times of high oxidative stress they can be associated with mitochondria as well as with membrane-less foci called stress granules (Miller-Fleming *et al*. 2014; Bankapalli *et al*. 2015). While *HSP32* and *HSP33* have almost complete sequence homology, *HSP31* moderately differs in sequence and performs additional activities compared to its paralogs. The second reason for focusing on Hsp31 was because it is the yeast ortholog of the human DJ-1 protein (PARK7) that is associated with early-onset autosomal recessive Parkinson’s disease(Bonifati *et al*. 2003). DJ-1 has been reported to have many cellular functions in the context of neuronal health (Bankapalli *et al*. 2015; Aslam and Hazbun 2016). Whether some of these functions contribute to neuronal death remain unresolved (Tsai *et al*. 2015; Mazza *et al*. 2022; Susarla *et al*. 2023).

In our study, we found that *HSP31* is selectively upregulated under conditions of mitochondrial protein import clogging combined with Mfb1 loss, but not in either condition alone. This indicates a specific transcriptional response to severe mPOS. Hsp31 may act as a holdase, preventing protein aggregation and/or facilitating the degradation of unimported mitochondrial proteins in the cytosol. This idea is supported by the finding that disruption of *HSP31* severely reduces the growth of *aac2^A128P, A137D^* and *aac2^A128P, A137D^ mfb1Δ* cells and increases ubiquitinated protein levels in the aggregates formed within these mutant cells. As Hsp31-GFP remains largely cytosolic under clogging conditions, Hsp31 likely defends the cytosol against mPOS. Interestingly, we also observed that global protein ubiquitination is significantly increased even in respiring *hsp31Δ* cells. This further suggests a protective role of Hsp31 against proteostatic stress and mPOS in cells that have a greater requirement for mitochondrial biogenesis and oxidative metabolism.

Approximately 10% of Parkinson’s disease (PD) cases are associated with mutations in single genes including human genes Fbxo7 (or PARK15) and DJ-1 (Shojaee *et al*. 2008; Di Fonzo *et al*. 2009). Fbxo7 is a mitochondria-associated F-box protein. It is known to act either as a substrate adaptor protein in SCF-type E3 ubiquitin ligase complex that regulates proteasomal and other cellular activities, or to function in a SCF-independent manner to promote mitophagy (Nelson *et al*. 2013; Joseph *et al*. 2018; Kraus *et al*. 2023). Interestingly, the SCF-independent function of Fbxo7 is required to stabilize Tomm20, the mammalian homolog of Tom20 (Teixeira *et al*. 2016). Furthermore, loss of Fbxo7 sensitizes cells to proteostatic stress in mitochondria (Kraus *et al*. 2023). Fbxo7 and Mfb1 share similarities only in the highly conserved F-box motif. Whether Fbxo7 plays a role in the maintenance of mitochondrial protein import competency remains to be tested. This function could become more relevant in actively respiring cells and under pathophysiological conditions with increased mitochondrial protein import stress. On the other hand, Hsp31/DJ-1 may be critical for protecting the cytosol against proteotoxicity caused by unimported mitochondrial proteins. Hence, both Mfb1 and Hsp31 may work in concert at the mitochondrial surface or in the cytosol, respectively, to support cellular proteostasis. Our data therefore draw mechanistic parallels between yeast Mfb1 and human F-box proteins such as Fbxo7. Additionally, our finding that HSP31 responds to mitochondrial import stress aligns with DJ-1’s broader role in cellular defense mechanisms, potentially providing a functional link between mitochondrial protein import stress, proteostasis, and neurodegeneration.

Altogether, our study reveals a role for Mfb1 in the maintenance of mitochondrial protein import competency. Furthermore, our work clarifies that Mfb1 likely functions in the absence of a SCF or non-SCF E3 ligase complex. It is important to appreciate that our work utilizes an endogenous mitochondrial protein (Aac2) which when mutated causes protein import to partially clog. Thus, discovering an anti-mPOS function for Mfb1 in this cellular setting is clinically relevant. Lastly, our work establishes a link between the disruption of Mfb1 during clogging and the activation of the heat shock protein Hsp31. In summary, we define a novel role for Mfb1 in protecting against mitochondrial precursor overaccumulation stress, and show that Hsp31 acts as a downstream proteostatic buffer in Mfb1-deficient cells. This reveals a cooperative axis between mitochondrial surface quality control (Mfb1) and cytosolic chaperone response (Hsp31). Studying this link further may help improve our understanding of how mitochondrial damage, protein import stress and cytosolic proteostasis are intricately connected in the pathogenesis of Parkinson’s disease.

## Experimental procedures

### Yeast strains and growth media

The genotypes and sources of yeast strains used in this study are listed in Supplemental Table 4. The *aac2^A128P^-mfb1Δ* interaction was studied in the BY4741/4742 strain background, whereas experiments involving the *aac2^A128P, A137D^* allele were carried out in strains of W303-1B background because expression of this severe clogger allele causes cell lethality in BY4741/4742 strains. Yeast cells were cultured in either YPD (1% yeast extract, 2% peptone and 2% dextrose), YPGR (1% yeast extract, 2% peptone, 2% galactose and 2% raffinose), YPGE (1% yeast extract, 2% peptone, 2% glycerol and 2% ethanol), minimal medium (0.67% yeast nitrogenous base without amino acids, 2% dextrose, supplemented with auxotrophic requirements), minimal medium with reduced glucose (0.67% yeast nitrogenous base without amino acids, 0.5% dextrose, supplemented with auxotrophic requirements), and sporulation media (0.5% yeast extract, 1% peptone, 0.05% glucose). All reagents, unless otherwise indicated, were obtained from Sigma-Aldrich.

### Gene disruption

The null alleles for *TOM5, TOM6, TOM7, UBX2, MFB1* and *HSP31* marked by *kan*, were transferred from strains of the yeast knockout collection into the W303-1B background by PCR amplification and one-step gene replacement. For disruption of *AMK1 (or YNL146C-A)*, a *URA3* cassette was used to replace the gene coding region. Correct disruption of the genes was confirmed by PCR, using primers listed in Supplemental Table 4.

### Mitochondrial isolation

Depending on the downstream application, mitochondria were isolated using either a glass-bead or Dounce homogenizer protocol. For mini-TurboID proximity ligation and determination of protein ubiquitination levels, 150 ml of yeast cultures were harvested at 5,000 RPM (J-10 rotor) for 10 minutes, washed once with water, and resuspended in 800 μl of SHP buffer (0.6 M sorbitol, 20 mM HEPES-KOH, PMSF, protease inhibitors, 20 mM NEM). Glass beads (Sigma-Aldrich) were added to the samples, before being vortexed twice for 20 seconds each, and maintained on ice. Samples were spun at 800 x g for 2 minutes to remove cell debris and the glass beads. The supernatant was spun at 12,000 x g for 15 minutes. The resulting supernatant was discarded, and the pellet was resuspended in the miniTurboID lysis buffer (50 mM Tris-HCl pH7.4, 150 mM NaCl, 1 mM EGTA, 1.5 mM MgCl_2_, 0.4% SDS and 1% NP-40) and stored at −80°C.

For TMT-MS, 400 ml of yeast cultures grown overnight at 25°C in YPGE were harvested at 5000 RPM (J-10 rotor) for 10 minutes and washed once with water. This was followed by washes with fresh TD buffer (100 mM Tris-SO_4_, pH9.4, 10 mM DTT) and SP buffer (1.2 M sorbitol, 20 mM potassium phosphate, pH7.4) buffers. Next, the samples were incubated in 5 mg/ml zymolyase prepared in SP buffer and rotated at 25°C for 2 hours to digest the cell wall. Following this, the samples were washed again with SP buffer and then resuspended in 20 ml of SHP buffer (0.6 M sorbitol, 20 mM HEPES-KOH, pH7.4, PMSF, protease inhibitors). Samples were placed in a Dounce homogenizer one at a time and exposed to 12-14 strokes to mechanically disrupt the cell membrane. Samples were spun at 2500xg (J-20 rotor) for 5 minutes to collect any unbroken spheroplasts and the supernatant spun at 12000xg for 10 minutes to pellet mitochondria. The supernatant containing “unclean” cytosol was spun again at 13000xg for 30 minutes to remove any membranous debris and the supernatant was designated as “clean” cytosol to be used for TMT-MS.

### MiniTurboID and mass spectrometry

#### MFB1-MiniTurboID biotin ligase fusion strains: growth, harvest, affinity purification

MiniTurbo biotin ligase ORF was fused to the C-terminal end of the *MFB1* gene immediately before the stop codon in the yeast chromosome. This fusion gene was generated in the WT and *aac2^A128P, A137D^* background strains. The resulting strains were referred to as WTmT and DMmT, respectively. These strains, as well as untagged wild type (WT) and *aac2^A128P, A137D^* (DM), were inoculated onto minimal medium supplemented with auxotrophic amino acids and allowed to grow overnight at 30°C. The strains were then inoculated into 150 ml of a non-fermentable medium (YPGE with additional tryptophan) and allowed to grow for 12 hours. Each strain was grown in triplicates. Crude mitochondria were isolated from these strains using the glass-bead protocol described above (see ‘Mitochondrial Isolation’). Bradford assay was performed to quantify mitochondrial yield. For each strain, 600-750 μg mitochondrial lysate was used as input for incubation with 300 μl of 50% slurry of Streptactin-Sepharose beads (IBA Lifesciences). Prior to incubation, the beads were equilibrated by washing twice with lysis buffer (50 mM Tris-HCl, pH7.4, 150 mM NaCl, 1 mM EGTA, 1.5 mM MgCl2, 0.4% SDS and 1% NP-40). The total volume of input lysate and beads in the lysis buffer was calculated such that lysate would bind the beads at a concentration between 1 – 1,25 μg/μl. Each sample was placed on a rotating drum at 4°C for 2 hours. After this, samples were spun (1000xg, 1 minute) to collect the unbound fraction. The beads were washed twice using wash buffer (1x buffer W) and the washes were kept as controls. After the wash steps, the beads were incubated with elution buffer (1x BXT), rotated using a Nutator mixer at room temperature for 10 minutes, and then spun (1000xg, 1 minute). The elution step was repeated once more. After confirming the presence of bait protein in the eluates, the elution fractions were combined and sent for mass spectrometry to obtain label-free quantification of the eluted proteins in the samples.

#### Sample digestion and cleanup

Fifty μg of proteins were digested using a modification of the FASP method(Wisniewski *et al*. 2009). In-solution proteins were denatured with SDS (1% final concentration), reduced and alkylated with TCEP and chloroacetamide (10 and 40 mM final concentration) at 70°C for 5 minutes. After cooling, the proteins were added to a 10 kDa MWCO membrane filter (Pall, OD010C34) with 200 µL of solution “UA”: 8 M urea with 100 mM tris pH 8.5. The filters were centrifuged at 14,000 x *g* until nearly dry, then rinse three times with 100 µL of UA solution and three times with 100 µL of 50 mM ammonium bicarbonate. The proteins were digested overnight at 37°C using 0.5 µg of trypsin in 75 µL of 50 mM ammonium bicarbonate. The resulting peptides were recovered from the filtrate by adding 50 µL of 1% TFA and centrifuging across the filter, followed by desalting on 2-core MCX stage tips (3M, 2241)(Rappsilber *et al*. 2003). The stage tips were activated with ACN followed by 3% ACN with 0.1% TFA. Next, samples were applied, followed by two washes with 3% ACN with 0.1% TFA, and one wash with 65% ACN with 0.1% TFA. Peptides were eluted with 75 µL of 65% ACN with 5% NH4OH (Millipore, 5.33003), and dried.

#### LC-MS methods

Samples were dissolved to a concentration of 0.25 µg/µL in water containing 2% ACN and 0.5% formic acid. Two µL were injected onto a pulled tip nano-LC column (New Objective, FS360-75-10-N) with 75 µm inner diameter packed to 25 cm with 3 µm, 120 Å, C18AQ particles (Dr. Maisch, r13.aq.0001). The column was maintained at 50°C with a column oven (Sonation GmbH, PRSO-V2). The peptides were separated using a 120-minute gradient from 3 – 28% ACN, followed by a 7 min ramp to 85% ACN and a 3 min hold at 85% ACN. The column was connected inline with an Orbitrap Lumos (Thermo) via a nanoelectrospray source operating at 2.5 kV. The mass spectrometer was operated in data-dependent top speed mode with a cycle time of 2.5s. MS1 scans were collected at 120000 resolution with AGC target of 6.0E5 and maximum injection time of 50 ms. CID fragmentation was used followed by MS2 scans in the ion trap with AGC target 2.0E3 and 38 ms maximum injection time.

#### Database searching and label-free quantification

The MS data was searched using SequestHT in Proteome Discoverer (version 2.4, Thermo Scientific) against the S. cerevisiae proteome from Uniprot, containing 6816 sequences and a list of common laboratory contaminant proteins. Enzyme specificity was fully tryptic with up to 2 missed cleavages. Precursor and product ion mass tolerances were 10 ppm and 0.6 Da, respectively. Cysteine carbamidomethylation was set as a fixed modification. Methionine oxidation, lysine biotinylatin, protein-terminal methionine loss and acetylation were set as a variable modification. The output was filtered using the Percolator algorithm with strict FDR set to 0.01. Label-free quantification was performed in Proteome Discoverer.

### Cytosolic fraction quantitative mass spectrometry

#### Post-Mitochondrial Cytosolic Fraction Preparation

TMT-MS was performed on the following yeast strains: WT*, mfb1Δ, aac2^A128P, A137D^* and *aac2^A128P, A137D^ mfb1Δ*. Each strain was represented in triplicates. Mitochondria were isolated using the ‘Dounce Homogenizer’ protocol as described above and removed from each sample using differential centrifugation to obtain the “mitochondria-free” cytosolic fraction (see above in the ‘Mitochondrial isolation’ section). The cytosolic fractions were quantified using the Bradford Assay, resuspended at a final concentration of 1ug/uL in 1xSHP Buffer (0.6M sorbitol, 20mM HEPES KOH, PMSF, and protease inhibitors). Each sample was submitted at a final volume of 1mL in Eppendorf tubes on ice to the SUNY Upstate proteomics core.

#### Sample digestion, labeling and cleanup

Samples received by the proteomics core were prepared for multiplexed quantitative mass spectrometry. Samples were buffer exchanged on a 3 kDa molecular weight cutoff filter (Amicon 3k Ultracel) using 50 mM triethylammonium bicarbonate (Thermo). One hundred µg was taken for digestion using an EasyPep Mini MS sample prep kit (Thermo, A40006). To each buffer-exchanged sample, 65 µL of lysis buffer was added followed by 50 µL of reduction solution and 50 µL of alkylating solution. Samples were incubated at 95°C for 10 minutes, then cooled to room temperature. To each sample 4 µg of trypsin / Lys-C protease was added and the reaction was incubated at 37°C overnight. TMT reagents were reconstituted with 20 µL acetonitrile (ACN) and the contents of each label added to a digested sample. After 60 min, 50 µL of quenching solution was added, consisting of 20% formic acid and 5% hydroxylamine (v/v) in water. The labeled digests were cleaned up by a solid-phase extraction device contained in the EasyPep kit, and dried by speed-vac. The individually labeled samples were dissolved in in 100 µL of 3% ACN and 0.2% trifluoroacetic acid (v/v) in water, and 15 µL of each was used to create a pooled sample consisting of 180 µg.

#### Fractionation

Following an LC-MS experiment to check digestion and labeling quality of the pooled samples, 100 µg of the pooled sample was fractionated using a Pierce High pH Reversed-Phase Peptide Fractionation Kit (part # 84868), per the manufacturer’s instructions for TMT-labeled peptides. In brief, samples were dissolved in 300 µL of 0.1% trifluoroacetic acid in water, and applied to the conditioned resin. Samples were washed first with water and then with 300 µL of 5% ACN, 0.1% triethylamine (TEA) in water. The second wash was collected for analysis. Peptides were step eluted from the resin using 300 µL of solvent consisting of 5 to 50% ACN with 0.1% TEA in eight steps. All collected fractions were dried in a speed-vac.

#### LC-MS/MS

Dried fractions were reconstituted in 25 µL of load solvent consisting of 3% ACN and 0.5% formic acid in water, and a 5 µL aliquot was diluted with 7 µL of the same solvent. Of these 12 µL, 2 µL were injected onto a pulled tip nano-LC column (New Objective, FS360-75-10-N) with 75 µm inner diameter packed to 28 cm with 2.4 µm, 120 Å, C18AQ particles (Dr. Maisch). The column was maintained at 50°C with a column oven (Sonation GmbH, PRSO-V2). The peptides were separated using a 135-minute gradient consisting of 3 – 12.5% ACN over 60 min, 12.5 – 28% over 60 min, 28 - 85 % ACN over 7 min, a 3 min hold, and 5 min re-equilibration at 3% ACN. The column was connected inline with an Orbitrap Lumos (Thermo) via a nanoelectrospray source operating at 2.5 kV. The mass spectrometer was operated in data-dependent top speed mode with a cycle time of 3s. MS^1^ scans were collected from 375 – 1500 m/z at 120,000 resolution and a maximum injection time of 50 ms. HCD fragmentation at 40% collision energy was used followed by MS^2^ scans in the Orbitrap at 50,000 resolution with a 125 ms maximum injection time.

### Yeast transcriptomic analysis

#### Total RNA extraction

For RNA-seq analysis, three biologically independent replicates were used for each strain. Yeast strains were first grown in a minimal medium supplemented with auxotrophic amino acids at 30°C overnight (50 ml). They were then removed from the minimal medium, inoculated into YPGE (100 ml culture), and grown at 25°C until they reached an OD600 between 1 and 2. Cells were then harvested and placed immediately on ice followed by a modified chloroform-methanol-based RNA extraction protocol(Schmitt *et al*. 1990). In this protocol, the cells were suspended in RNA extraction buffer (0.1M NaCl, 10 mM EDTA, 5% SDS, 50 mM Tris-HCl), diluted with equal volumes of PCI (25:24:1 mixture of phenol:chloroform:isoamyl alcohol with phenol at a pH of 4.5) and allowed to sit at room temperature for 5 minutes. The samples were then incubated with glass beads and subjected to a bead-beater apparatus twice for 30 seconds with a 30-second interval on ice. The homogenized samples were centrifuged (13,000 RPM for 10 minutes) and the aqueous phase was obtained. The aqueous phase was subjected to extraction twice again with PCI and once with 100% chloroform. The final aqueous phase was mixed with 10% volume of 3 M sodium acetate pH 5.2 and 2 volumes of 95% ethanol and the sample allowed to precipitate at −20°C for one hour. The pellet was washed with 70% ethanol, resuspended with DEPC-treated water, and stored at −80C before being submitted for RNA sequencing.

#### cDNA Library preparation and RNA Sequencing

RNA concentration was confirmed by NanoDrop spectrophotometer, and Agilent Bioanalyzer was used for determining RNA integrity. Sequencing libraries were prepared from samples with the Illumina Stranded mRNA Library Prep kit, using one microgram of RNA as input. The standard protocol was followed and 10 cycles of PCR amplification were. Sequencing was performed on a NextSeq2000 instrument using a paired end 2×100bp read. For sequencing, a P3 sequencing kit was used which had 1.362B reads pass filter. Approximately ∼28M reads per sample were obtained.

#### RNAseq Analysis and Data Visualization

Raw reads and counts generated were processed using the Partek software. The processed dataset containing differentially expressed gene lists was analyzed in R studio using the DESeq2 package (Love *et al*. 2014). For stratified analysis, the DESeq model was generated using two groups of samples at a time to allow for a 1-1 comparison. For multifactorial analysis, the DESeq model was generated using all four groups of samples to identify additive genetic interaction effects.

### Triton insoluble fraction extraction

A previously described procedure was adapted for extracting triton-insoluble proteins (Coyne *et al*. 2023a). Briefly, cells were resuspended in Triton insoluble lysis buffer (0.5% Triton X-100, 150 mM NaCl, 50 mM HEPES-KOH pH 7.4, 1 mM EDTA with PMSF and protease inhibitors) followed by vigorous bead beating (two rounds of 20 seconds vortexing in the presence of glass beads) and placed on ice. Samples were spun at a low speed (600xg) to collect debris, unbroken cells, etc. The supernatant was placed in a fresh tube and subjected to a 20000xg spin for 30 minutes. After discarding the supernatant, the pellet was washed with lysis buffer and spun once more (20000xg for 30 minutes). The final pellet was designated as “triton-insoluble” and was resuspended using 8M urea and 5% SDS prior to western blot analysis.

### Western blot analyses

Protein lysates were run on SDS-PAGE gels, and proteins were transferred overnight to PVDF membranes. Total protein stain (LICORBio) was performed for all membranes, prior to blocking (5% milk in 1xTBST) and downstream antibody incubation steps. For miniTurboID-MS, the PVDF membranes were probed with either anti-V5 (Cell Signaling cat#E9H80) or anti-Biotin (Milipore Sigma cat#B3640) antibody. For TMT-MS, the PVDF membranes were probed with custom-made or commercially available primary antibodies targeting Porin, Abf2, Aco1, Aac2, or Pgk1, followed by the appropriate secondary antibody (goat anti-rabbit or goat anti-mouse). For the miniTurboID-MS experiment, membranes from the biotinylated protein elution assay were probed with either anti-Biotin (Milipore Sigma cat#B3640) or anti-V5 (Cell Signaling cat#E9H80) antibodies before being probed with Donkey anti-goat secondary, or Goat anti-Mouse secondary antibody, respectively. Finally, for the global ubiquitination measurement, membranes were probed with an anti-ubiquitin (Invitrogen cat#10H4L21) primary antibody before being probed with Goat anti-Rabbit secondary antibody. Each secondary antibody was HRP-conjugated, which allowed for chemiluminescent detection of protein bands. For complete list of reagents, see Supplemental Table 5.

### Fluorescence Microscopy

Strains expressing Hsp31-mNG (chromosomally integrated) were grown in YPGE (1% yeast extract, 2% peptone, 2% glycerol, 2% ethanol) overnight, subcultured for 4 hours in fresh media until mid-log phase (OD_600_ between 0.5-1) and then prepared for imaging. Strains expressing Mfb1-GFP (*URA3*-based multi-copy plasmid) were grown overnight in selective media lacking uracil (YNBCasD+Adenine+Tryptophan), subcultured in YPGR until mid-log phase (OD_600_ between 0.5-1) and then prepared for imaging. Both Hsp31-GFP and Mfb1-GFP expressing cells were stained with MitoTrackerRed CMXROS Dye (final concentration 250 nM) for an hour in the dark, washed twice with sterile water, and then resuspended in fresh media prior to mounting on a glass slide. Cells were inspected and imaged using a 100× oil (NA 1.4) objective on a Zeiss Imager.Z1 fluorescence microscope with a Hamamatsu CCD camera. AxioVision software was used to capture and save images. The imaging filter sets included DIC, GFP, and TexasRed which were used for visualizing the general cellular morphology, GFP-fused protein, and mitochondrial profiles respectively. Scale bars were added using FIJI by utilizing the known distance-to-pixels ratio.

### Data availability

The RNA-Seq data generated as part of this study are presented as supplemental tables. The data have also been deposited to NCBI Gene Expression Omnibus/Sequence Read Archive with the accession numbers GSE298360 (reviewer access token: ahajikesbnuppgz). The TMT-MS and label-free quantitative proteomic data have been deposited to PRoteomics IDEntifications Database (PRIDE) with the accession numbers PXD064566 (reviewer access token: eRhPMf5huNXd) and PXD065210 (reviewer access token: V2mgzhjkPAff). All software used in this study is publicly available and there was no code generated as part of this study.

## Supporting information

This article contains supporting information.

## Acknowledgments

We thank Dr. Bruce Knutson, Dr. Ryan Palumbo, and Dr. Ebbing de Jong for help in the MiniTurboID assay and the analysis of proteomic data, Maya Schuldiner for sharing GFP-tagged yeast strains, and Victor Z. Chen for help in RNA-sequencing analysis. We acknowledge the use of the AI tool ChatGPT in assisting us with writing code for computational analysis in RStudio and for suggesting grammar edits to improve the clarity of our manuscript.

## Author contributions

**GM**: Conceptualization, Methodology, Formal Analysis, Investigation, Visualization, Data Curation, Writing – Original Draft and Review/Editing, Funding Acquisition. **XW**: Investigation, Data Curation, Validation, Resources, Methodology. **EF**: Investigation, Visualization. **AG**: consultation and supervision for computational analysis of RNAseq data, manuscript revision. **XJC**: Conceptualization, Investigation, Data Curation, Visualization, Formal Analysis, Methodology, Supervision, Writing – Original and Review/Editing, Funding Acquisition.

