## Supporting information for "The Mitochondrial F-box protein 1 and DJ-1 homolog HSP31 support cellular proteostasis during mitochondrial protein import clogging"

**The PDF file includes:**
Figure S1 to S7

**Other Supplementary Information for this manuscript include the following:**
Table S1 to S6

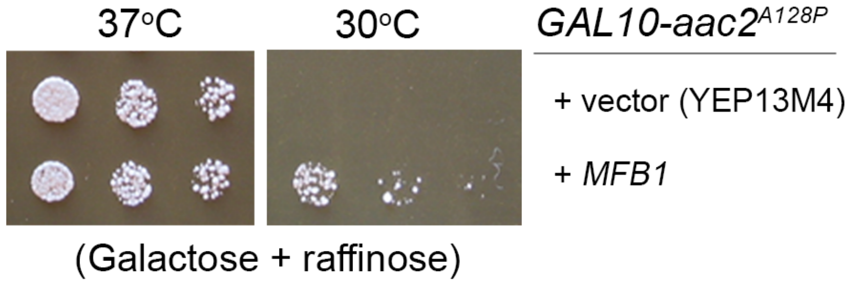


**Figure S1: A subclone containing only the *MFB1* open reading frame can rescue cell growth defect of a yeast strain expressing *aac2*$^{A128P}$.** The open reading frame of *MFB1* including its endogenous promoter was cloned into the multicopy YEP13M4 vector. This plasmid rescued the growth of cells expressing *GAL10-aac2*$^{A128P}$ at 30ºC on complete galactose and raffinose medium. The growth of *GAL10-aac2*$^{A128P}$ cells is suppressed at 37ºC due to global repression of protein synthesis, which is used as a control for the experiment.

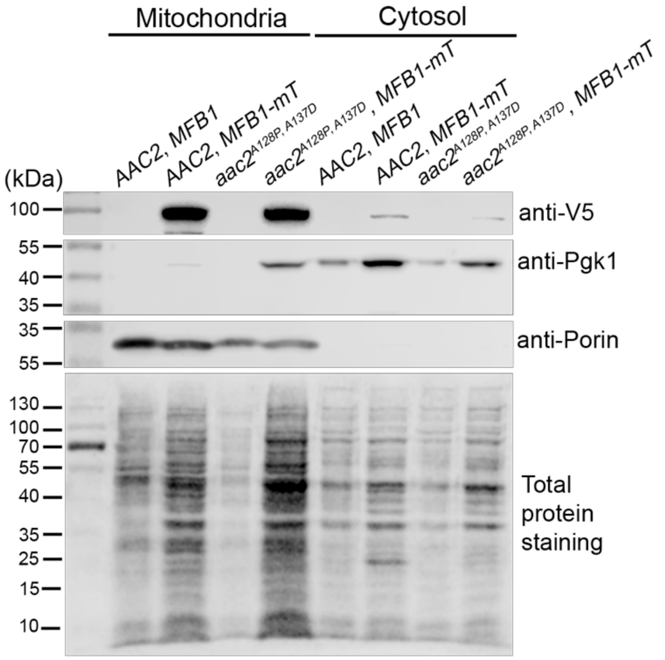


**Figure S2: Western blot showing the localization of the Mfb1-miniTurboID fusion protein to the mitochondrial fraction**. Yeast cells were first cultured on minimal medium before being inoculated in the complete medium with the non-fermentable ethanol plus glycerol as carbon sources. Mitochondrial and cytosolic fractions were prepared and analyzed by SDS-PAGE followed by western blot. Anti-V5 antibody was used to monitor the subcellular distribution of the Mfb1-miniTurboID fusion protein that is tagged with the V5 epitope. Antibodies against Pgk1 and porin were used as markers for cytosolic and mitochondrial fractions, respectively.

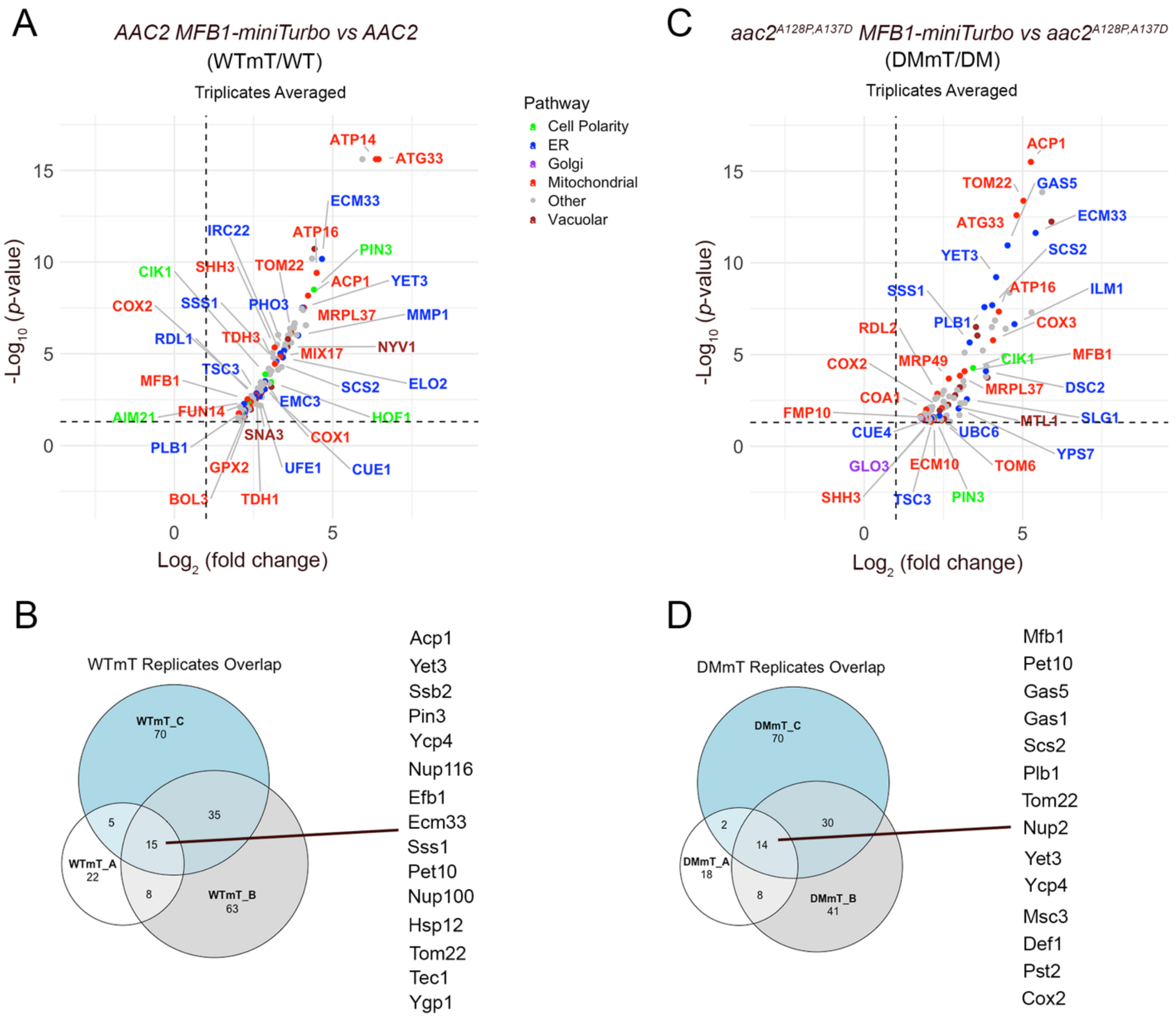


**Figure S3: Mfb1's interactome as revealed by BioID assay. (A)** Volcano plot showing biotinylated proteins in the Mfb1-miniTurboID tagged cells in the background of wild type AAC2 (WTmT), normalized to untagged wild type after averaging three biologically independent replicates. **(B)** Venn diagram of proteins common across the triplicates described in (A). **(C)** Volcano plot showing biotinylated proteins in the Mfb1-miniTurboID tagged cells in the clogger aac2A128P, A137D cells (DMmT), normalized to untagged clogger after averaging three biologically independent replicates. **(D)** Venn diagram of proteins common across the triplicates described in (C).

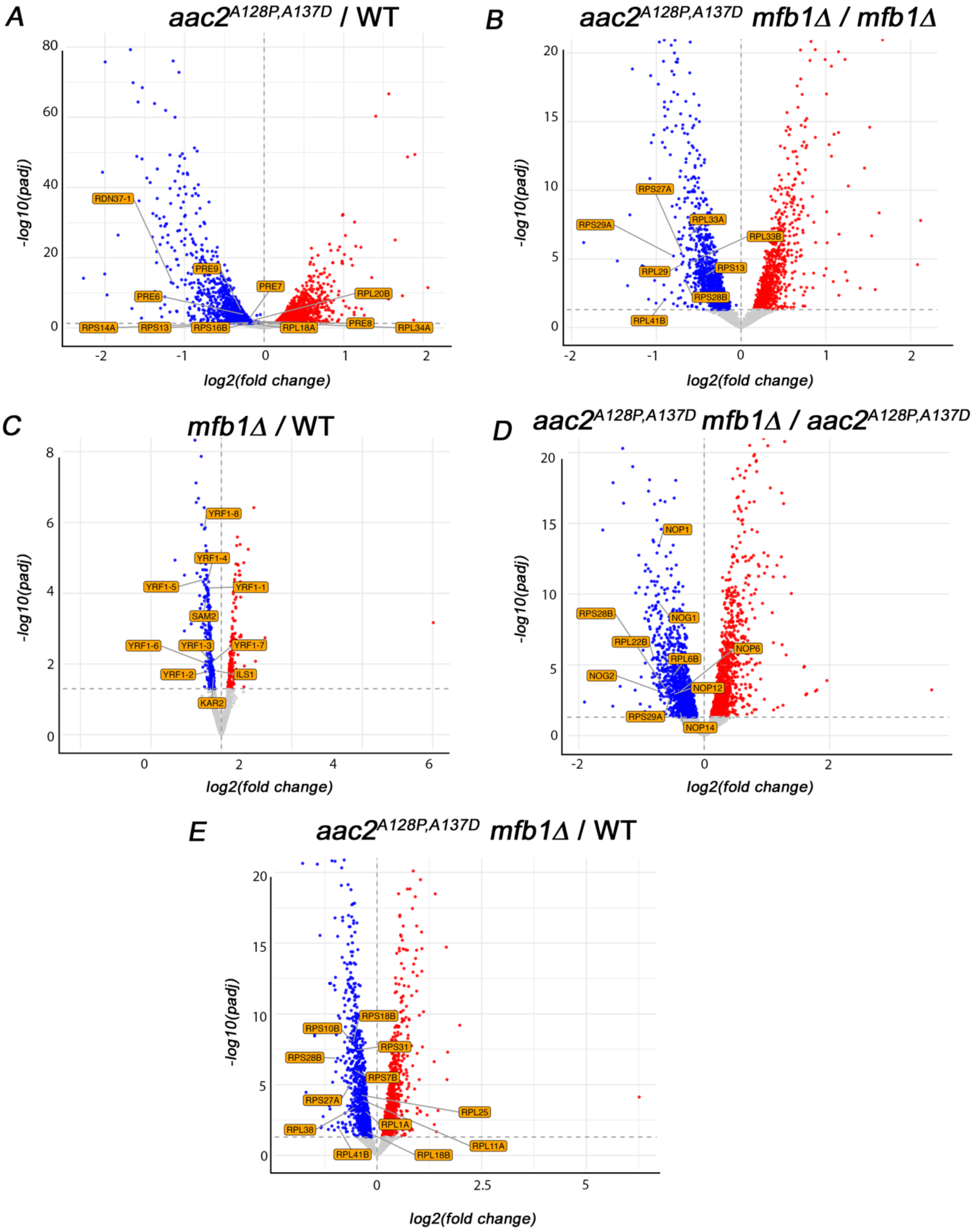


**Figure S4: Downregulated genes in the RNAseq dataset**. Volcano plots showing differential gene expression highlighting downregulated genes in the stratified analysis when comparing *aac2*$^{A128P, A137D}$ vs WT **(A)**, *aac2*$^{A128P, A137D}$ *mfb1Δ* vs *mfb1Δ* **(B)**, *mfb1Δ* vs WT **(C)**, *aac2*$^{A128P, A137D}$ *mfb1Δ* vs *aac2*$^{A128P, A137D}$ **(D)**, and *aac2*$^{A128P, A137D}$ *mfb1Δ* vs WT **(E)**. Significant genes have *FDR* (*padj*) <0.05 and absolute ($log_2$(fold change)) > 0.

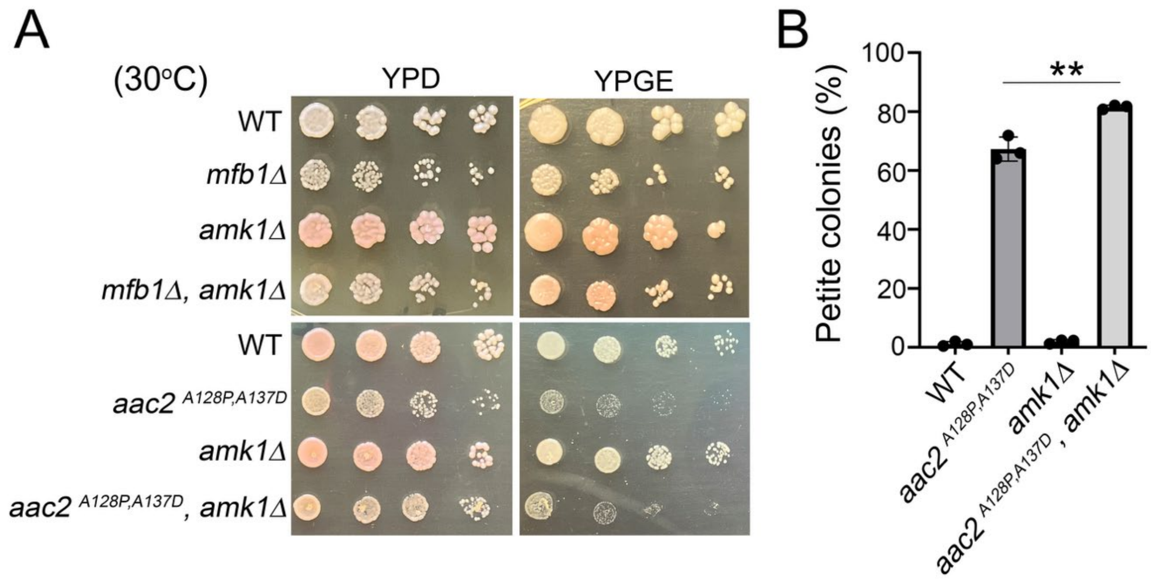


**Figure S5: Characterization of AMK1**. **(A)** Five-fold dilution series for yeast strains lacking either *MFB1* or *AMK1* under the AAC2 wild type or clogger (*aac2*$^{A128P, A137D}$) background. Ank1 loss does not exacerbate growth defects under clogging conditions for cells grown in either fermentable (YPD) or non-fermentable (YPGE) media. Note that Amk1-deficient strains have slightly greater red pigment accumulation when compared to wild type cells, suggesting a potential role for Amk1 in redox homeostasis. **(B)** Amk1 loss midly increases petite frequency under import clogging conditions. *P* value was calculated using Student's t-test. **, p<0.01.

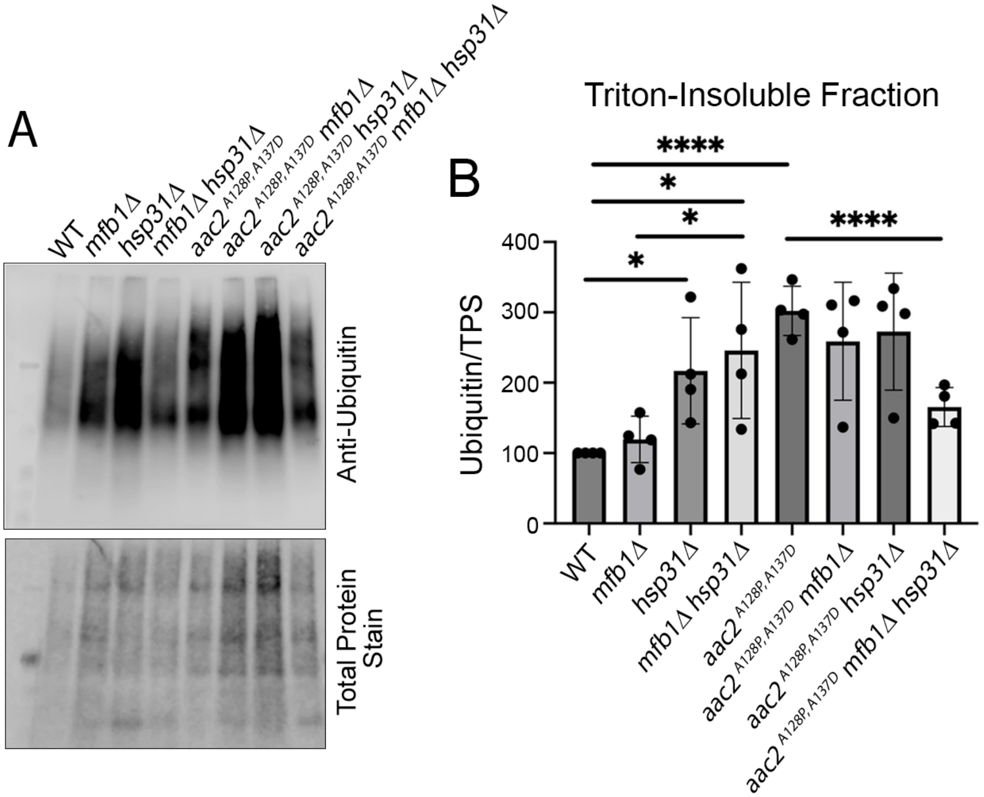


**Figure S6: Disruption of *HSP31* is sufficient to increase levels of ubiquitinated proteins in Triton-insoluble fractions. (A)** Western blot analysis of ubiquitinated proteins sequestered within insoluble aggregates. Samples were obtained from yeast strains grown under respiring conditions. **(B)** Quantification of western blot shown in (A). *P* values were calculated using Student's t-test. *, p <0.05; ****, p<0.0001.

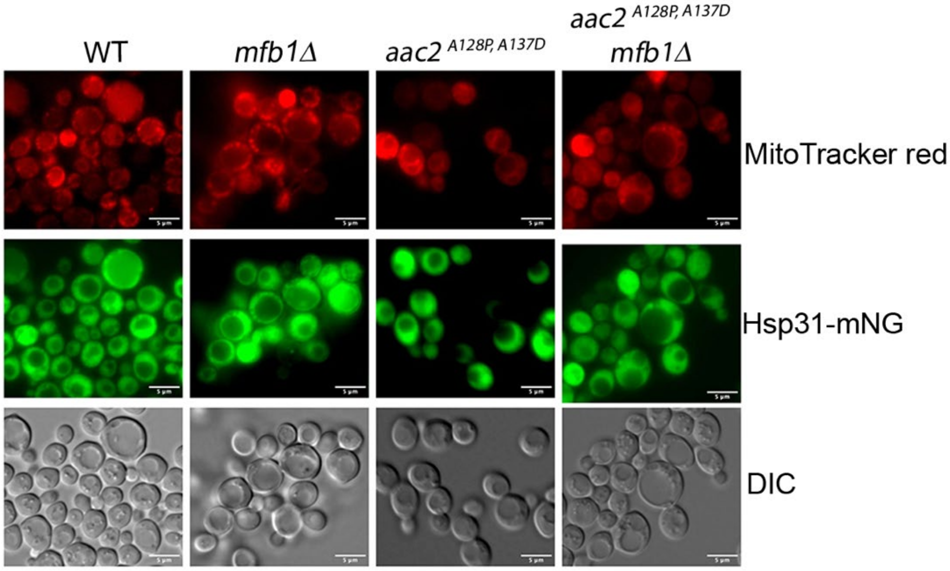


**Figure S7: Hsp31 remains cytosolic, regardless of clogging conditions or Mfb1-deficiency.** Representative images showing Hsp31-mNG fusion protein in the cytosol of all yeast strains visualized. Mitochondria were stained with Mitotracker Red.

Legends for Supplemental Tables.

**Supplemental Table 1:** Excel workbook containing pathway analysis results from Funspec for *aac2*$^{A128P,\ A137D}$ (DM) vs WT comparison. Gene Ontology (GO) terms and Munich Information Center for Protein Sequences (MIPS) classification terms are included. Sheet 1 contains upregulated pathway lists after Bonferroni correction ($p < 0.01$). Sheet 2 contains upregulated pathway lists without Bonferroni correction. Sheet 3 contains downregulated pathway lists after Bonferroni correction ($p < 0.01$). Sheet 4 contains downregulated pathway lists without Bonferroni correction.

**Supplemental Table 2:** Excel workbook containing pathway analysis results from Funspec for *aac2*$^{A128P,\ A137D}$*mfb1Δ* vs *aac2*$^{A128P,\ A137D}$ comparison. Gene Ontology (GO) terms and Munich Information Center for Protein Sequences (MIPS) classification terms are included. Sheet 1 contains upregulated pathway lists after Bonferroni correction ($p < 0.01$). Sheet 2 contains upregulated pathway lists without Bonferroni correction. Sheet 3 contains downregulated pathway lists after Bonferroni correction ($p < 0.01$). Sheet 4 contains downregulated pathway lists without Bonferroni correction.

**Supplemental Table 3:** Excel workbook containing pathway analysis results from Funspec for *aac2*$^{A128P,\ A137D}$*mfb1Δ* vs WT comparison. Gene Ontology (GO) terms and Munich Information Center for Protein Sequences (MIPS) classification terms are included. Sheet 1 contains upregulated pathway lists after Bonferroni correction ($p < 0.01$). Sheet 2 contains upregulated pathway lists without Bonferroni correction. Sheet 3 contains downregulated pathway lists after Bonferroni correction ($p < 0.01$). Sheet 4 contains downregulated pathway lists without Bonferroni correction.

**Supplemental Table 4:** Excel sheet listing the genes and their strength of genetic interaction given the genotype-environment multifactorial analysis of the RNAseq dataset. Each gene's expression level in response to *MFB1* disruption is tested in the *AAC2* wild type and *aac2*$^{A128P,\ A137D}$ mutant backgrounds. Strength of interaction is the difference in log2 of the fold change in the mutant vs wild type AAC2 backgrounds.

**Supplemental Table 5:** Excel workbook containing yeast strains (Sheet 1), plasmids (Sheet 2), and primers (Sheet 3) used in this study.

**Supplemental Table 6:** List of relevant reagents and sources used in this study.
